# Broad and thematic remodeling of the surface glycoproteome on isogenic cells transformed with driving proliferative oncogenes

**DOI:** 10.1101/808139

**Authors:** Kevin K. Leung, Gary M. Wilson, Lisa L. Kirkemo, Nicholas M. Riley, Joshua J. Coon, James A. Wells

## Abstract

The cell surface proteome, the surfaceome, is the interface for engaging the extracellular space in normal and cancer cells. Here we apply quantitative proteomics of N-linked glycoproteins to reveal how a collection of some 700 surface proteins is dramatically remodeled in an isogenic breast epithelial cell line stably expressing any of six of the most prominent proliferative oncogenes, including the receptor tyrosine kinases, EGFR and HER2, and downstream signaling partners such as KRAS, BRAF, MEK and AKT. We find that each oncogene has somewhat different surfaceomes but the functions of these proteins are harmonized by common biological themes including up-regulation of nutrient transporters, down-regulation of adhesion molecules and tumor suppressing phosphatases, and alteration in immune modulators. Addition of a potent MEK inhibitor that blocks MAPK signaling brings each oncogene-induced surfaceome back to a common state reflecting their strong dependence on the MAPK pathway to propagate signaling. Using a recently developed glyco-proteomics method of activated ion electron transfer dissociation (AI-ETD) we found massive oncogene-induced changes in 142 N-linked glycans and differential increases in complex hybrid glycans especially for KRAS and HER2 oncogenes. Overall, these studies provide a broad systems level view of how specific driver oncogenes remodel the surface glycoproteome in a cell autologous fashion, and suggest possible surface targets, and combinations thereof, for drug and biomarker discovery.

**Significant statement:** The cell surface glycoproteome (surfaceome) mediates interactions between the cell and the extracellular environment, and is a major target for immunotherapy in cancer. Using state-of-the-art proteomics methods, we compared how six neighboring proliferative oncogenes cause large and bidirectional expression of some 700 surface proteins and the 142 different glycans that decorate them. While each oncogene induces large and somewhat unique glycoproteomes relative to non-transformed cells, we find common functional consequences that are massively reversed by small molecule inhibition of the MAPK pathway. This large-scale comparative study provides important insights for how oncogenes remodel isogenic cells in a cell autologous fashion, and suggest possible new opportunities for antibody drug discovery in more complex tumor settings.

## Introduction

The cell surface proteome, or surfaceome, is the main interface for cellular signaling, nutrient homeostasis, cellular adhesion, and defines immunologic identity. To survive, cancer cells adjust to promote increased nutrient import, pro-growth signaling, and evasion of immunological surveillance among others (Hanahan and Weinberg, 2011). There are some 4,000 different membrane proteins encoded in the human genome (Gonzalez et al., 2010; Wallin and von Heijne, 1998), yet antibodies to only about two dozen cell surface targets have been approved for therapeutic intervention, prompting the need to discover novel tumor specific antigens (Carter and Lazar, 2018; Chames et al., 2009).

Recent surfaceome studies in an isogenic MCF10A breast epithelial cell line transformed with oncogenic KRAS have identified more than two dozen up-regulated surface proteins that function in a cell autologous fashion to promote increased cell proliferation, metastasis, metabolic activity and immunologic suppression (Martinko et al., 2018; Ye et al., 2016; Ying et al., 2012). Many of the most powerful oncogenes are linked to KRAS and the MAPK pathway including overactivation of receptor tyrosine kinases, such as EGFR and HER2, or mutations in BRAF or RAS (**Fig 1A**). It is well-known these neighboring oncogenes are mutually exclusive in human tumors. For example, lung cancer patients with oncogenic EGFR mutations seldom harbor oncogenic KRAS or BRAF mutations and vice versa (Cancer Genome Atlas Research Network et al., 2014). Also, recent evidence show that oncogene co-expression can induce synthetic lethality or oncogene-induced senescence further reinforcing that activation of one oncogene without the others is preferable in cancer (Cisowski et al., 2015; Unni et al., 2015).

**Figure 1:**
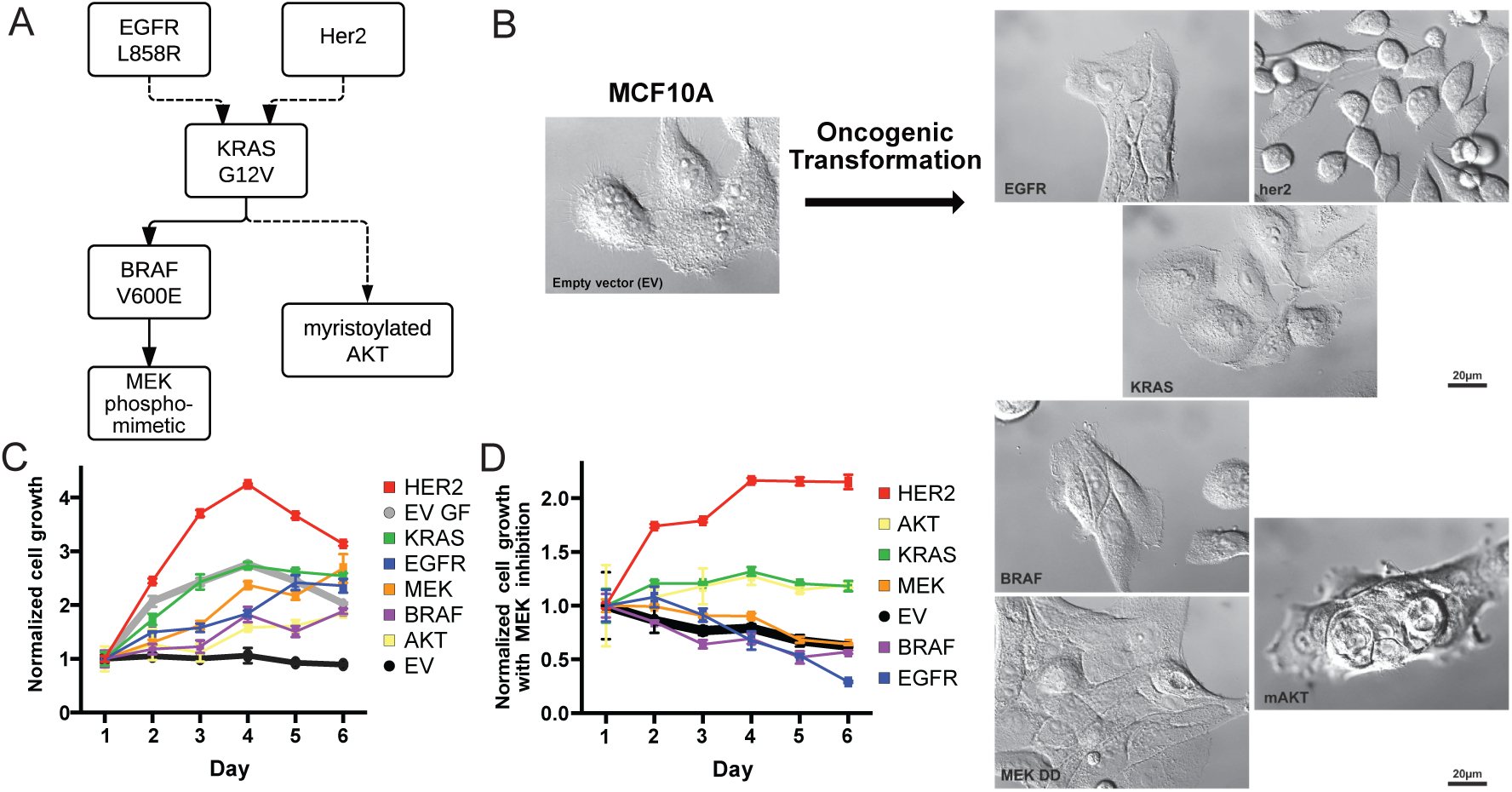
Growth rates and morphologies for MCF10A cells transformed with neighboring proliferative oncogenes that control the MAPK pathway. (A) Simplified signaling schematic relationship of six proliferative oncogenes studied, EGFR^L858R^, HER2 over-expression, KRAS^G12V^, BRAF^V600E^, MEK ^DD^ and AKT^myr^. (B) Oncogenic transformation of MCF10A induces diverse cellular morphologies. Note, images are presented in the same order of the schematic in panel A. (C) MCF10A cells stably transformed with lenti-virus with the different onocogenes grow independent of growth factor. Grey and black lines indicate cellular growth of MCF10A EV control with and without growth factors, respectively. Cell growth (n = 3) was measured each day for 6 days by CellTiter-Glo luminescent cell viability assay and normalized to viability on day 1. (D) Suppression of growth for all cell lines by treatment with 100 nM MEK inhibitor (PD0325901) in the absence of growth factors. MCF10A cells transformed with HER2 appears to be least sensitive to MEKi, followed by KRAS^G12V^ and AKT^myr^.

There is also substantial evidence that cell surface glycosylation is altered in cancer. Incomplete or truncated synthesis, extended branching, core fucosylation, and sialylation of cell surface glycans are hallmarks of tumor cells, which alter physiological mechanisms of cell-cell adhesion, communication, and immune system recognition (Fernandes et al., 1991; Hakomori, 1996; Hudak et al., 2014; Xia et al., 2016; Xiao et al., 2016; Kannagi et al., 2010; Varki and Gagneux, 2012). Over the past decade, chemical glycoproteomics has revealed specific examples of altered glycosylation and heterogeneity of glycans on particular proteins (Palaniappan and Bertozzi, 2016). However, we do not know how expression of different oncogenes globally alters glycosylation on individual proteins at a proteome-wide scale. Very recently, hybrid type ETD methods, such as activated ion electron transfer dissociation (AI-ETD) and ETD with supplemental activation (EThcD), have emerged as forerunners for glycoproteomic profiling due to their ability to generate sequence informative tandem mass spectra that links both the peptide backbone and corresponding glycan modification (Liu et al., 2017; Riley et al., 2019; Yang et al., 2017). These techniques can provide granularity to the altered glycosylation states of cell surface proteins upon oncogenic transformation.

Here we address how neighboring driver oncogenes (KRAS^G12V^, HER2 overexpression, EGFR^L858R^, BRAF^V600E^, phosphomemetic MEK^S218D/S222D^, and myristoylated AKT) stably expressed in a common non-cancerous isogenic epithelial cell alters the surface glycoproteome. Using cell-surface capture (CSC)(Bausch-Fluck et al., 2015; Wollscheid et al., 2009) and AI-ETD glycoproteomics (Riley et al., 2019), we found each oncogene induced common and unique sets of up and down-regulated surface proteins and associated glycans. These sets of protein revealed common biological themes including increased expression of nutrient transporters and decreased expression of adhesion molecules. These effects were massively reversed upon inhibition of the MAPK pathway emphasizing its central importance in remodeling the surfaceome. These oncogene-induced surface proteins highlight targets or combinations to consider for immunotherapy.

## Results

### Phenotypic analysis of oncogene transformed MCF10A cells

Tumor biology is highly complex and varies depending on the cell type of origin, stage, oncogenes, epigenome, stroma, vascularization, the immune system and metabolism. We deliberately took a simplistic reductionist approach to compare how specific proliferative oncogenes alter the cell surface proteome in a cell autologous fashion by using isogenic cells stably transformed with different oncogenes. No single cell type, culturing condition, or context is representative of all or even one cancer type. For practical reasons we chose the spontaneously immortalized breast epithelial MCF10A as our parental cell line because it is often used as a neutral starting point for oncogene studies (Allen-Petersen et al., 2014; Martins et al., 2015; Qu et al., 2015; Soule et al., 1990). Although MCF10A certainly does not recapitulate the diversity of mammary cell biology, it is of epithelial origin like most common tumors; it is non tumorigenic, requires growth factors for survival and does not harbor gene amplifications or other chromosomal aberrations typical of advanced cancer cell lines. Our intent is to compare how neighboring driving oncogenes remodel their surface glycoproteomes in isogenic cells in a cell autologous fashion.

Lentivirus was used to stably transform MCF10A cells with six prevalent oncogenes that are neighbors in proliferative signaling: HER2 overexpression, EGFR^L858R^, KRAS^G12V^, BRAF^V600E^, the phosphomimetic MEK^S218D/S222D^ (MEK^DD^) and myristoylated AKT (AKT^myr^) (**Figure 1A**). Remarkably, the morphologies of each of the transformed cells varied from each other when cultured in the absence of growth factors indicative of differences in proteomic landscape (**Figure 1B**). In the absence of growth factors, cells transformed with HER2 and KRAS^G12V^ grew to confluence, while cells harboring EGFR^L858R^, BRAF^V600E^, and MEK^DD^ did not reach confluency, indicative of contact dependent growth inhibition. Cells transformed with AKT^myr^, which signals through a parallel pathway relative to the other five oncogenes, had the most dramatic morphology change, displaying vertically stacked clusters of cells. Unlike the parental cell line, all the MCF10A cells stably transformed with any of the six oncogenes proliferated in the absence of growth factors to various degrees (**Figure 1C**). The HER2 and KRAS^G12V^ transformed cells proliferated most rapidly in the absence of growth factors, and even comparable or faster to the untransformed MCF10A cultured in the presence of growth factors. The HER2 and KRAS^G12V^ cells also lifted off the plates much more readily than the others, suggesting reduced adhesion phenotype.

These oncogenes can drive multiple branched pathways yet it was previously shown that inhibition of the MAPK pathway with the potent and selective MEK inhibitor (PD032590, MEKi) significantly reverses the surfaceome changes of MCF10A cells transformed with KRAS^G12V^ (Martinko et al., 2018). Indeed, MEKi substantially hampered growth for all cell lines either in the absence or presence of growth factors (**Figure 1D**, **Figure S1**). Overexpression of HER2 was most resistant to MEKi, followed by KRAS^G12V^ and AKT^myr^, whereas cells containing EGFR^L858R^, BRAF^V600E^, or MEK^DD^ were most sensitive to MEKi.

### Differential expression of oncogene-induced surfaceomes in MCF10A cells

We next probed how the cell surfaceome is altered in the oncogene transformed cells compared to the empty vector control. N-glycosylation is present on >85% of cell surface proteins (Apweiler et al., 1999) and can be exploited to capture the N-glycosylated proteins using a biotin hydrazide enrichment method called cell surface capture (CSC) (Bausch-Fluck et al., 2018; Wollscheid et al., 2009). Here, we utilized a CSC protocol coupled with stable isotope labeling by amino acids in cell culture (SILAC) to compare the surfaceomes from the oncogene transformed MCF10A cells to the empty vector control (**Figure S2**) (Leung et al., 2019). We identified and quantified a total of 654 cell surface proteins across the six oncogenic cell lines (**Figure 2A**). Remarkably, the expression for 43% of the aggregate surface proteins (280 of 654) was altered by at least two-fold for the oncogene transformed cell lines relative to empty vector control reflecting significant remodeling of the surfaceomes. In each of the six datasets, we observed at least two-fold changes for 100 to 150 different surface proteins (**Figure 2A, Figure S3A-F**); these changes were evenly split between up or down regulated sets reflecting bi-directional remodeling. Many of the differentially expressed proteins overlapped, but each cell line had a substantial number of uniquely differentially regulated proteins, presumably resulting from slight differences in signaling between each oncogene. Although it is well known that correlations between protein and RNA levels are not often strongly correlated (Lundberg et al., 2010; Martinko et al., 2018; Schwanhäusser et al., 2011), we were prompted to determine whether the most common changes we observed. BCAM and NRCAM downregulation, in particular, were also observed in patient samples harboring the same oncogenic signatures in a number of cancers types of epithelial origin (**Figure 2B-C**). Utilizing 17 TCGA provisional dataset from The Cancer Genome Atlas with carcinoma (epithelial origin) annotation (Cancer Genome Atlas Research Network et al., 2013), activating oncogenic signature was defined as G12 or Q61 mutation in KRAS, L858 or amplification of EGFR, V600E mutation in BRAF, or amplification of HER2. Transcriptional upregulation of MME, however, was not found in any of the dataset searched, suggesting regulation at the translational level or additional factors at play.

**Figure 2:**
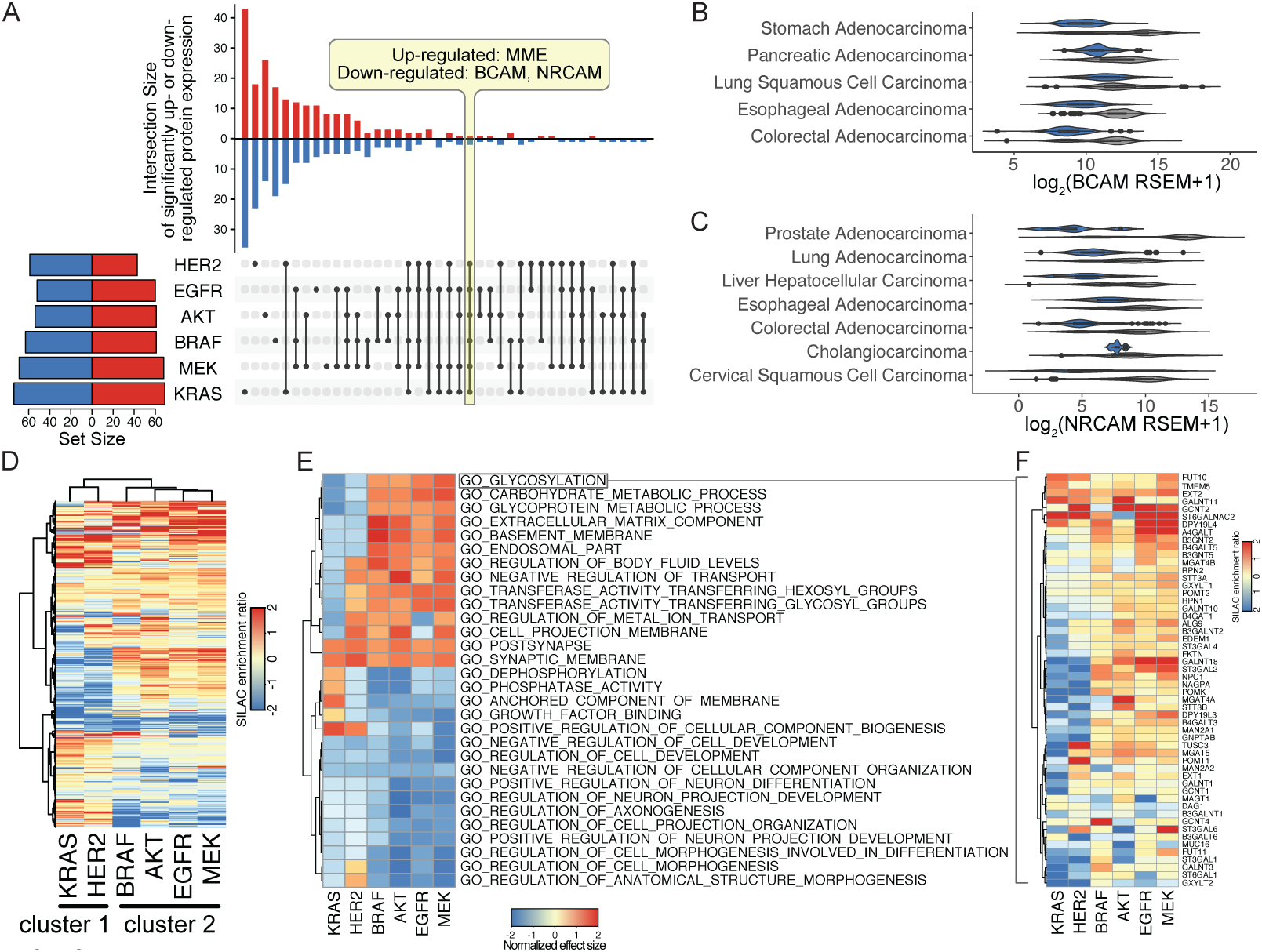
Proliferative oncogenes cause large changes in the surfaceome that are diverse in detail but have common functional themes. (A) Many differentially regulated proteins are unique to each cell line, and only three proteins were commonly up or down-regulated among all six cell lines. In the top bar graph, up-regulated proteins (red) are indicated by the upward bars, and down-regulated proteins (blue) are indicated by the downward bars. The specific overlapping groups are indicated by the black solid points below the bar graph. Total differentially regulated proteins for each cell line are indicated in the left bar graph. Up-regulated (red) and down-regulated (blue) proteins are defined as log_2_(FC) > 1 and p value < 0.05. (B-C) BCAM (B) and NRCAM (C) are significantly down regulated across various carcinoma with activating oncogenic signature in KRAS, HER2, BRAF and EGFR (blue) compared to no mutations in these genes (grey). Activating oncogenic signature was defined as G12 or Q61 mutation in KRAS, L858 or amplification of EGFR, V600E mutation in BRAF, or amplification of HER2. Genomic and expression data was obtained from 17 TCGA provisional dataset with carcinoma (epithelial origin) annotation. (D) Hierarchal clustering of surfaceome changes revealed similarities between HER2 and KRAS transformed cells. (E) Top 30 enriched gene sets identified by Gene Set Enrichment Analysis (GSEA) of the proteomics dataset using Gene Ontology terms shows clustering between HER2 and KRAS transformed cells. Positive normalized effect size (up-regulation) is shown in red and negative (down-regulated) normalized effect size is shown in blue. Proteins were pre-ranked by median SILAC peptide ratio and GSEA was performed using MySigDB C5 GO gene set collection. (F) Proteins involved in glycosylation (GO:0070085) are down regulated in the KRAS and HER2 cluster (blue) while the same proteins are up-regulated in the other oncogenes cluster. Heatmap for each Gene Set identified is also available at wellslab.ucsf.edu/oncogene_surfaceome.

At a global level, there were greater similarities between particular oncogenes. For example, cells harboring KRAS^G12V^ and HER2 clustered more closely together (Cluster 1), and those containing BRAF^V600E^, AKT^myr^, EGFR^L858R^ and MEK^DD^ clustered together (Cluster 2) as seen either in the upset plot (**Figure 2A**) or a heatmap with hierarchical clustering (**Figure 2D**). The Pearson correlation coefficients were also higher within rather than across the two Clusters (**Figure S4A**). One of the striking findings was the upregulation of HER2 expression in the KRAS^G12V^ transformed cells but not in Cluster 2 transformed cells (**Figure S5**), which may help to explain the stronger similarity in oncogene-induced surfaceomes between the HER2 and KRAS^G12V^ cell lines. This same analysis also showed striking compensating regulation, where HER2 is down-regulated in the EGFR oncogene expressing cell line.

Despite detailed differences at the individual target level, these harmonized into common biological processes when viewed by Gene Set Enrichment Analysis (GSEA) (**Figure 2E**). For example, glycosylation and carbohydrate metabolism were similarly altered features for all the oncogenes (**Figure 2F**), consistent with numerous reports that altered glycosylation correlates with the development and progression of cancer (Pinho and Reis, 2015; Varki et al., 2015). In addition, we see strong down-regulation of proteins involved in differentiation and adhesion reflecting cell attachment and migration (**Supplemental Fig 6A**). There were also large changes in some cell surface phosphatases involved in down-regulation of receptor tyrosine kinases (**Figure S6B)**. These specific heatmaps reinforce the general division between Cluster 1 and Cluster 2. Due to the complexity of this datatype, a data browser was made to view each individual gene sets identified (https://wellslab.ucsf.edu/oncogene_surfaceome).

### MEK inhibition induces common surfaceome changes

The MAPK pathway is a central driver of cell proliferation and has been a major therapeutic target in cancer. Indeed, MEKi significantly impedes the growth of all the oncogene transformed cells either in the absence (**Figure 1D**) or presence of growth factors (**Figure S2**). To determine how MEKi alters the surfaceome of the oncogene transformed cells, we compared the proteomic landscape of each oncogenic cell line in the presence and absence of MEKi. The proteomics dataset identified and quantified a total of 772 proteins, including an intersection of 492 proteins with the empty vector dataset discussed above, for a total of 934 proteins quantified between the two experiments. (**Figure 3A-C**).

**Figure 3:**
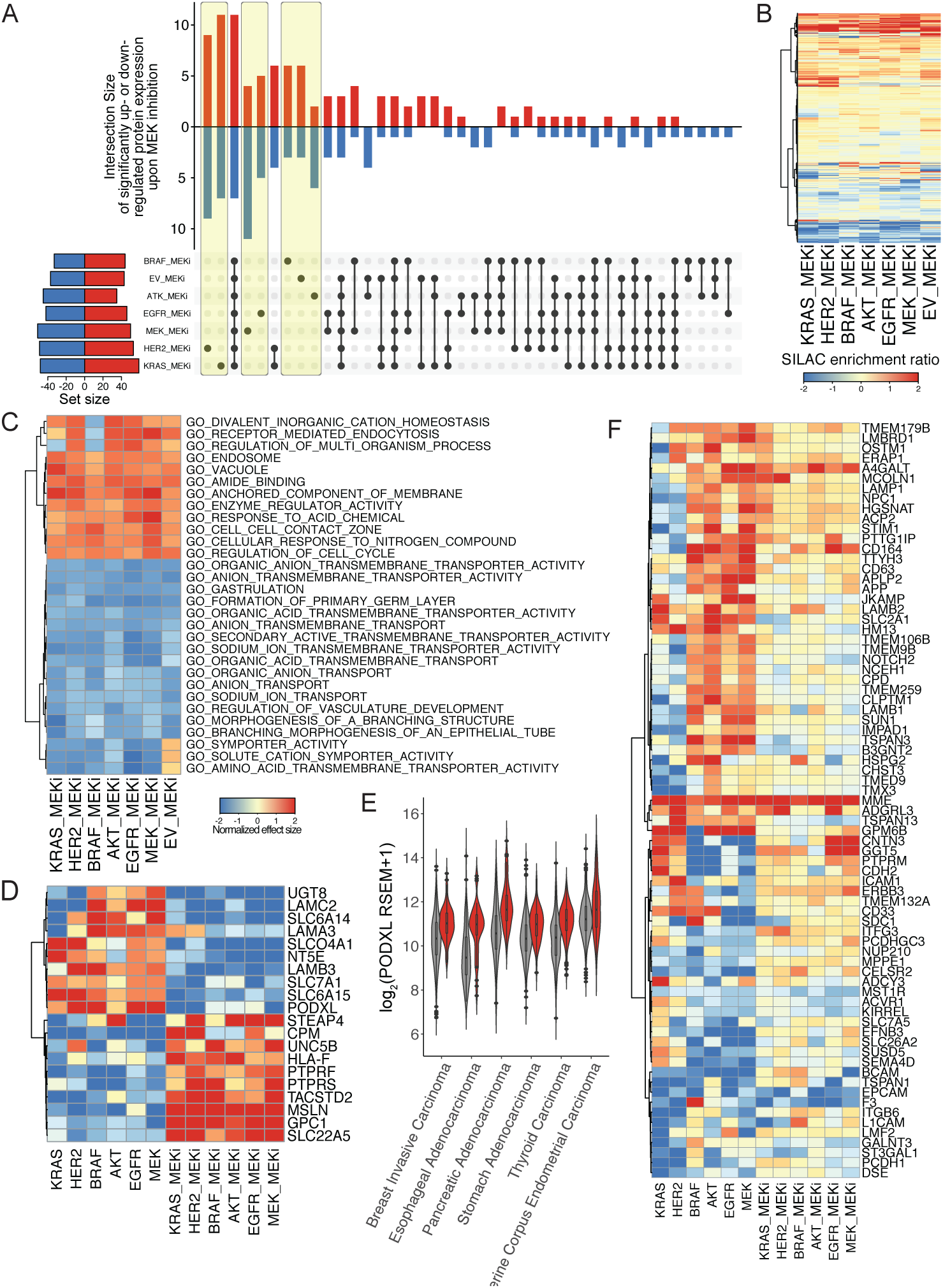
Inhibition of MEK led to similar perturbations of the surfaceome regardless of oncogenic transformation. (A) Majority of differentially regulated surface proteins were either completely unique (highlighted in yellow) or completely common (bar 3) to all seven cell lines. In the top bar graph, up-regulated genes (red) are indicated by the upward bars, and down-regulated genes (blue) are indicated by the downward bars. The specific overlapping groups are indicated by the black solid points below the bar graph. Total differentially regulated proteins for each cell line are indicated in the left bar graph. Up-regulated (red) and down-regulated (blue) proteins are defined as log_2_(FC) > 1 and p value < 0.05. (B) Hierarchal clustering of surfaceome changes shows MEK inhibition treatment affected each cell line similarly. (C) Top 30 enriched gene sets identified by Gene Set Enrichment Analysis (GSEA) of the proteomics dataset using Gene Ontology terms show similarities between all cell lines, indicating unique changes could be functionally redundant. Positive normalized effect size (up-regulation) is shown in red and negative (down-regulated) normalized effect size is shown in blue. Proteins were pre-ranked by median SILAC ratio and GSEA was performed using MySigDB C5 GO gene set collection. Heatmap for each Gene Set identified is also available at wellslab.ucsf.edu/oncogene_surfaceome. (D) MEK dependent regulation of protein expression induced by oncogenic transformation. Proteins shown were differentially regulated by oncogene (either log_2_(FC) > 1 or log_2_(FC) < -1 in oncogene vs EV) and were then reversed (flipped log_2_(FC) < -1 and log_2_(FC) > 1 in MEK inhibition vs respective cell line) in at least 3 cell lines. (E) PODXL is significantly upregulated across various carcinoma with activating oncogenic signature in KRAS, HER2, BRAF and EGFR (red) compared to no mutations in these genes (grey). Activating oncogenic signature was defined as G12 or Q61 mutation in KRAS, L858 or amplification of EGFR, V600E mutation in BRAF, or amplification of HER2. Genomic and expression data was obtained from 17 TCGA provisional dataset with carcinoma (epithelial origin) annotation. (F) MEK independent regulation of protein expression induced by oncogenic transformation. Proteins shown were differentially regulated by oncogene (either log_2_(FC) > 1 or log_2_(FC) < -1 in onco vs EV) and were then reverted to a median level but not reversed (flipped log_2_(FC) < -1 and log_2_(FC) > 1 in MEK inhibition vs respective cell line) in at least 3 cell lines.

In large measure MEKi reversed the effects of the oncogenes and remarkably induced a more common state between all the cell lines including the empty vector control. For example, in contrast to the uninhibited datasets where 43% of proteins were differentially regulated by more than two-fold in the oncogene expressing cells, only 17% of the proteins (129 of 772) were altered by more than two-fold in the presence of MEKi (**Figure 3A**). Of the proteins detected, 18 proteins were commonly changed across all oncogenes, as opposed to three in the absence of MEK inhibition (**Figure 3A, Figure S3G-M**). The similarity can be seen by hierarchical clustering of the MEKi datasets showing common changes across all cell lines (**Figure 3B**). The Pearson correlation between the six oncogene and untransformed control were also higher in general (**Figure S4B**). MEKi in KRAS^G12V^ and HER2 are still most closely correlated. GSEA of the MEKi data indicated a general common phenotypic reversal with down regulation of membrane transporters, metabolism, and upregulation of cell adhesion proteins consistent with a decrease in cancer associated phenotypes such as cellular proliferation and metastasis (**Figure 3C, Figure S6C-D**).

Integrating oncogenic transformation and MEK inhibition dataset, 20 protein targets symmetrically flip from being significantly up to down regulated or vice versa in at least three cell lines, suggesting that the expression of these proteins is strongly dependent on the MAPK signaling pathway (**Figure 3D**). One such target, PODXL, appears to be stringently regulated by the MAPK pathway. Utilizing the same informatics approaches described above, transcription of PODXL was also found to be strongly upregulated in several cancer types of epithelial origin (**Figure 3E**). Additionally, 75 targets that were markedly up or down regulated revert to a median level of expression upon treatment with MEKi in at least three cell lines (**Figure 3F**). Interestingly, there were a handful of oncogene-induced targets that are further up-regulated upon MEKi, such as MME, suggesting they are maintained by circuitous pathways outside of MAPK (MEKi independent). Overall, the six oncogenes cause profound changes to the surfaceome that alter common biological processes, which can be largely blunted by inhibition of the MAPK pathway.

### Oncogenes induce large changes to the glycoproteome

Glycosylation has long been known as a biomarker for cancer (Adamczyk et al., 2012; Mechref et al., 2012; Peracaula et al., 2008; Pinho and Reis, 2015) and our GSEA data shows systematic up regulation of proteins involved in glycosylation, especially in Cluster 2 (**Figure 2E**). We thus sought to identify the N-glycosylation modifications on specific membrane proteins for the six oncogene-transformed cell lines to compare among themselves and the empty vector control. For maximal coverage we enriched the glycoproteomes using the lectin Concanavalin A (ConA) and hydrophilic interaction liquid chromatography (HILIC), that provide a complementary means for capturing high mannose type and complex type glycans, respectively (Totten et al., 2017; Zhang et al., 2016). We processed the N-glycoproteomes in biological triplicate utilizing LC-MS/MS coupled with activated ion electron transfer dissociation (AI-ETD) (**Figure S7**) (Riley et al., 2017). AI-ETD fragmentation combines radical-driven dissociation and vibrational activation and was recently shown to afford robust fragmentation of intact glycopeptides (Riley et al., 2019). Spectral assignments were made using the Byonic search engine (Bern et al., 2012); glycopeptides that were not identified across each of the three biological replicates for each cell line were removed. (**Figure S8**)

Combining ConA and HILIC enrichments allowed for sampling of different glycan classes and the aggregated results of intact N-glycoproteomic analyses identified a total of 2,648 unique N-glycopeptides that passed conservative filtering described in Materials and Methods (**Fig 4, S9**). Of this total, 189 were redundant glycan-glycosite pairs, i.e., glycoforms that resulted from miscleaved or differentially oxidized peptides leaving a total of 2,459 unique glycoforms (**Figure 4A**). These glycoforms map to 785 N-glycosites found on a total of 480 glycoproteins. Thus, on average each glycoprotein had 1.6 N-linked glycosites and each site had on average 3.1 different glycans, reflecting significant heterogeneity in glycosylation. Similar, yet unique glycoforms indicated that the heterogeneity of protein N-glycosylation is driven largely by the biology and not artifacts of in-source fragmentation. Of the 785 glycosites, 324 are either known or predicted to be glycosylated in UniProt (UniProt Consortium, 2010), and over half are newly reported here.

**Figure 4:**
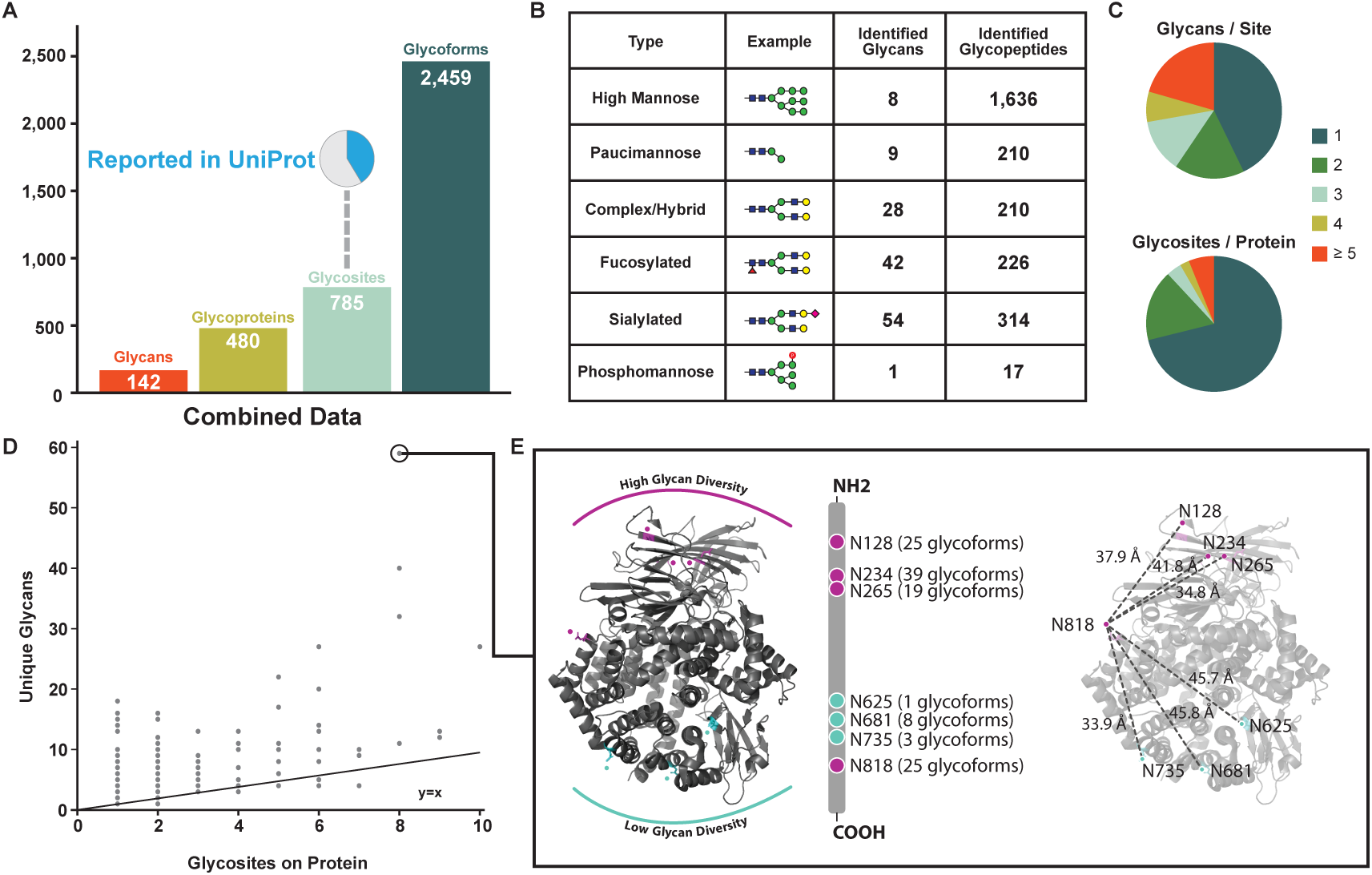
A global overview of the MCF10a glycoproteome identified from all cell lines. (A) Unique glycans, glycoproteins, glycosites, and glycoforms identified across all six cell lines. A pie charts represents a high percentage of glycosites not previously reported in uniport. (B) High mannose glycan type is the most common glycan identified. A total of six common glycan types are used to categorize glycans identified. (C) The varying degrees of microheterogeneity (top) and macroheterogeneity (bottom) of protein N-glycosylation. (D) A scatter plot of unique glycans versus number of glycosites for each protein provides an additional view of heterogeneity in protein N-glycosylation. The protein with the most heterogenous glycoform was Aminopeptidase N (ANPEP, UniProtKB: P15144). (E) Mapping glycan diversity on structure of ANPEP reveals spatial regulation of glycosylation patterns. Asparagine residues with high (magenta) and low (teal) numbers unique glycans are highlighted. The most C-terminus glyan, N818, has high glycan diversity distance and has similar distance to all other asparagine residues. (PDB ID: 4FYR)

In total we identified 142 different glycan structures. The glycans can be categorized into six structural classes based on their maturation state as they transition from ER to Golgi and then split off to either lysosomal, granular, or cell surface destinations (**Figure S10**)(Yarema and Bertozzi, 2001). The six glycan categories represent a gradation of maturation from the least mature high mannose, to paucimannose, phosphomannose, complex/hybrid, and finally fucosylated and sialylated (**Figure 4B**). We identified 1,636 mannose glycopeptides containing any of 8 high mannose-type glycans and 210 were found trimmed to contain any of 9 different paucimannose structures. The remaining 767 N-glycopeptides had more complex glycans including: 210 complex/hybrid glycopeptides with 28 different structures, 314 of the sialylated class with 54 different structures, and 226 of the fucosylated class with 42 different structures. We found only 17 phosphomannose containing glycopeptides each having the same structural type that are on proteins typically found in lysosomes (Kornfeld and Mellman, 1989; Rohn et al., 2000; Rouillé et al., 2000). Thus, over 60% of the N-glycopeptides identified contained less mature high mannose or paucimannose structures originating from processing in the ER or early Golgi. It is not surprising to find these high mannose modifications associated with cell surface proteins as others have seen that high mannose N-glycosylation is abundant at the cell surface and especially associated with oncogene transformation (An et al., 2012; Balog et al., 2012; Holst et al., 2016).

We found significant heterogeneity in the number of different glycan structures at any given site (**Figure 4C**). Approximately 45% of the sites were observed with only one glycan structure, but the range of glycans on these sites was broad. Some glycosites, such as position N-234 on aminopeptidase N (ANPEP) had up to 39 different glycans detected. Additionally, the number of glycosites per protein varies considerably; about 70% of the glycoproteins have only one site of N-glycosylation but about 10% have over five sites. There is a general trend between the number of glycosites identified on a given protein and the number of unique glycans it has; however, some proteins show significantly increased heterogeneity of glycans relative to their number of total glycosites (**Figure 4D**). The most extreme glycan diversity was observed on ANPEP, on which 59 unique glycans were identified across 8 glycosites (**Figure 4E**). Interestingly, the glycosites with higher glycan diversity mapped to specific regions on the 3D structure of ANPEP. One face of the protein containing a cluster of four asparagines (at positions 128, 234, 265 and 818) had high glycan diversity ranging from 19 to 39 different glycan structures per glycosite, while the opposite face containing three clustered asparagines (at positions 625, 681 and 735) had much lower diversity, ranging between 1 and 8 different glycan structures per site. This suggests topological bias of glycosylation on the folded protein structure.

We next applied quantitative analysis to compare the different oncogene transformed cell lines in terms of their glycoproteome landscapes and changes between them and the empty vector control. We observed hundreds of glycopeptides that were significantly and differentially expressed (q < 0.05) upon oncogenic transformation relative to the empty vector control (**Figure S11**). In general, the volcano plots were symmetric reflecting bi-directional changes and the fold-changes ranged from 50-fold down to 50-fold upregulated representing significant remodeling of the glycoproteome. Changes in glycosylation were greater than changes in surface protein expression which range from 2-8 fold (**Figure S3**). We quantified about 600 glycopeptides in each data set, of which two-thirds are shared in all data sets (**Figure 5A**). There was high quantitative reproducibility of the glycopeptide measurements based on the close hierarchical clustering of the three biological replicates (**Figure 5B**), tight clustering by principal component analysis, and low (<30%) median CV from the empty vector control cell line (**Figure S12A-B**). MEK^DD^ and BRAF^V600E^ had the greatest glycoproteome similarity, while empty vector was the furthest removed from all the oncogenes.

**Figure 5:**
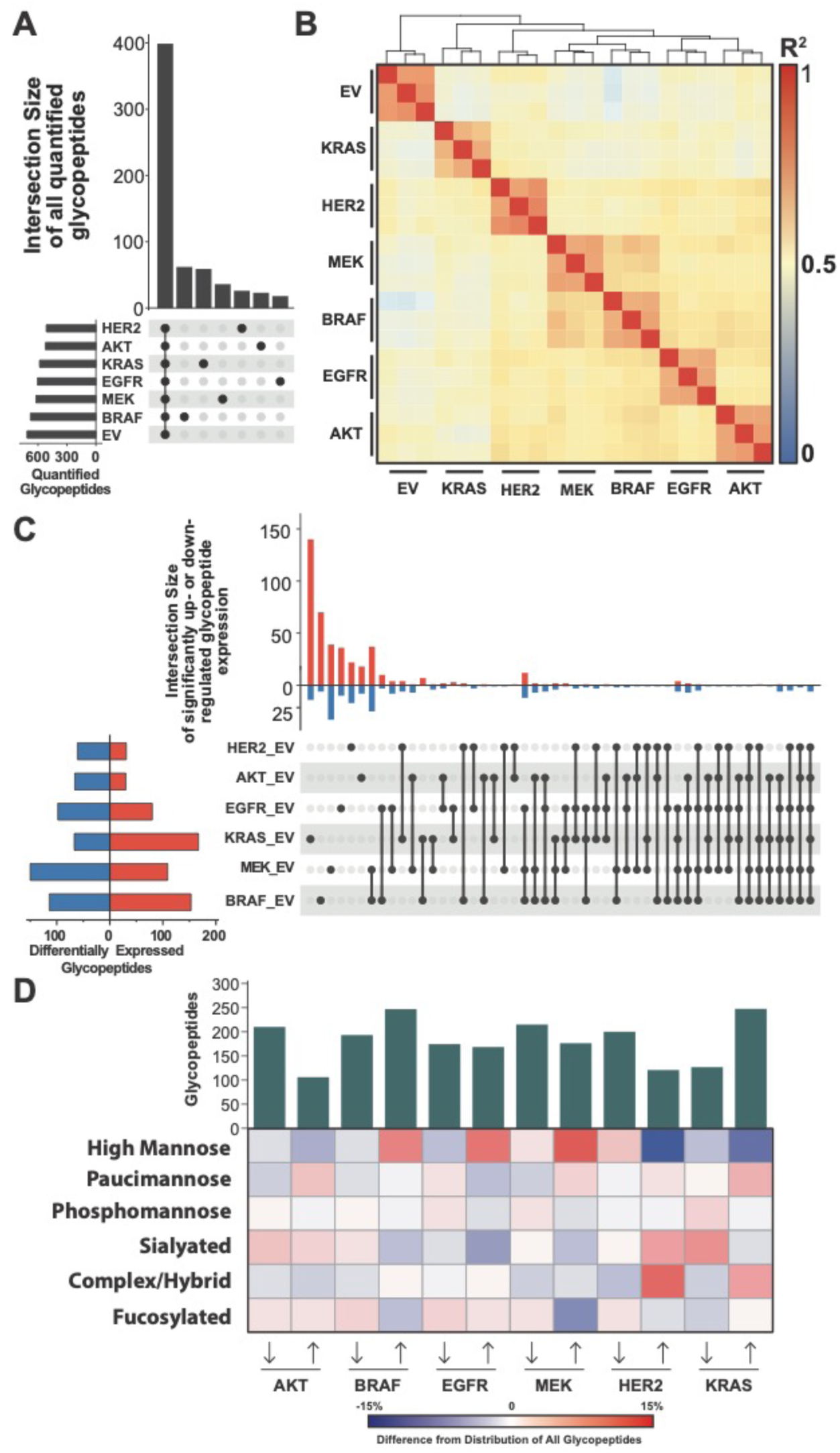
Quantitative glycopeptide measurements across mutant cell lines. (A) An UpSet plot shows glycopeptide identifications that are unique to or shared between data sets. (B) Pairwise Pearson correlations from all replicate analyses illustrates clustering between MEK^DD^ and BRAF^V600E^ glycoproteome. (C) An UpSet plot displays the shared and unique glycopeptides that are differentially expressed upon oncogenic transformation. (D) A heatmap display the differences glycan type distribution for up and down regulated glycopeptides across cell lines.

The UpSet plot in **Figure 5c** displays significant glycopeptide differential expression that is shared and unique to each cell line. MCF10A transformed with the KRAS^G12V^ oncogene resulted in the largest set of uniquely changing glycopeptides. 154 of the 234 differentially expressed glycopeptides in the KRAS^G12V^ cell line were unique to KRAS^G12V^ transformation. Some of these were highly protein specific. For example, 28 of the 154 glycopeptides uniquely differentially expressed by KRAS^G12V^ were identified from ANPEP and all were upregulated upon oncogenic transformation as was the protein itself (**Figure S13**). Other glycopeptides from ANPEP were differentially expressed in different sets of cell lines. In fact, 51 of 69 glycopeptides from ANPEP were differentially expressed in at least one cell line; 12 ANPEP glycopeptides were significantly upregulated in HER2 and 5 were shared between HER2 and KRAS^G12V^.

BRAF^V600E^, EGFR^L858R^, and MEK^DD^ shared the most overlap of significantly changing glycopeptides between any group of three cell lines (**Figure 5C**); 24 glycopeptides belonged to this intersection. 12 were overexpressed and 12 were under expressed upon oncogenic transformation. Laminin subunit alpha-3 (LAMA3) and N-acetylglucosamine-6-sulfatase (GNS) were most represented in the over expressed and under expressed group, respectively. Four glycopeptides from each protein were identified and significantly differentially expressed in these three cell lines. Six glycopeptides were significantly differentially expressed across all 6 cell lines, 3 of which were identified from galectin-3 binding protein (Gal-3BP) and all were downregulated glycopeptides modified with high mannose glycans (**Figure S14**).

We explored the glycan composition of differentially expressed glycopeptides to capture a broader view of differential glycosylation in the oncogene transformed cell lines and to see if general trends emerged. The heatmap in **Figure 5D** displays the differential glycome composition of glycopeptides changing more than 2-fold upon oncogenic transformation compared to empty vector control. We, again, observe greatest similarity between BRAF^V600E^, EGFR^L858R^, and MEK^DD^ cell lines, which have an increased proportion of high mannose glycans in upregulated glycopeptides. In contrast, HER2 and KRAS^G12V^ expressed fewer upregulated high mannose-modified glycopeptides and showed an increased proportion of complex/hybrid type glycopeptides. Further inspection revealed that nearly all the upregulated glycopeptides with a complex/hybrid glycan from the cell lines harboring HER2 (12 of 12) and KRAS^G12V^ (13 of 18) mapped to ANPEP. This protein was also upregulated on the KRAS^G12V^ surfaceome (Martinko et al., 2018), displayed the highest degree of glycan heterogeneity within the glycoproteomic data, and has previously been implicated in tumorigenesis (Dong et al., 2000; Pasqualini et al., 2000).

## Discussion

Oncogenesis is a complex phenomenon that involves aberrant changes in multiple biological processes to promote cancer cell survival (Hanahan and Weinberg, 2011). Here we study how the surfaceome remodels in a simplified cell autologous model by six prevalent and neighboring oncogenes that drive proliferation through the MAPK signaling node. Genetic studies have shown that these oncogenes typically exhibit mutual exclusivity in tumors from cancer patients (Cancer Genome Atlas Research Network et al., 2014). The surfaceome is a terminal manifestation of these signaling pathways. We find significant differences in detailed expression patterns consistent with differences in feedback loops and collateral signaling pathways between these oncogenes (Vaseva et al., 2018). However, we find these oncogene-induced surfaceome differences harmonize in similar functional outcomes overall, and consistent with observed mutual exclusivity.

We took a reductionist approach starting from an immortalized epithelial cell line stably transformed with each of the six different oncogenes to determine how the surfaceome is remodeled in a cell autologous fashion. This system is clearly an approximation and does not recapitulate the complexity of tumors that vary in cellular context, genetic variation, heterogeneity, oncogene expression, in the presence of host immune system and metabolism. Nonetheless, cell culture models are a practical reality allowing isogenic comparison between oncogenes, renewable access to materials that permits studies to be readily reproduced (Domcke et al., 2013; Rockwell, 1980; Wilding and Bodmer, 2014). We chose the spontaneously immortalized breast epithelial cell line, MCF10A, where the functional loss of p16^INK4a^ allow cells to be cultured with a myriad of oncogenes without oncogene-induced senescence. We picked this over cancer cell lines which are already transformed or other artificially immortalized cell lines because they often contain genetic lesions that may cause genomic instability, leading to a more idiosyncratic background. Overall, we find that independent expression of each of these six oncogenes induced profound changes in the surfaceome, both in the proteins expressed and the glycans that decorate them.

### Common phenotypes and biological themes induced by the six oncogenes

Although the six oncogenes we studied (HER2 overexpression, EGFR^L858R^, KRAS^G12V^, BRAF^V600E^, MEK^DD^, and AKT^myr^) have many variants that could have different phenotypes, we chose well known representatives to begin to understand the similarities and differences between them at a course-grain level. Transformation with each of these oncogenes led to rapid growth to varying degrees in the absence of growth factors and produced somewhat different cell morphologies characteristic of a cancer cell transformed phenotype. Each oncogene caused large changes in the surfaceome where about 40% of the detected N-glyco proteins had altered expression level by more than two-fold evenly split between up and down-regulated proteins reflecting bi-directional cell surface remodeling.

There were important differences between the oncogenes, and they clustered into two groups based on growth rates, surfaceomes and associated glycans. Cluster 1 containing HER2 overexpression and KRAS^G12V^ was most aggressive in proliferation and reduced adhesion. Cluster 2 was less aggressive included EGFR^L858R^, BRAF^V600E^, MEK^DD^ and AKT^myr^. We believe much of the similarity between HER2 and KRAS derives from KRAS induced expression of HER2, whereas EGFR suppresses expression of HER2 probably reflecting their signaling redundancy. MCF10A is derived from normal breast epithelial cells and among these oncogenes HER2 overexpression is most commonly seen in breast cancer patients at 13% compared to 2% for KRAS, 2.8 % for EGFR, 5% for AKT1, 1.7% for BRAF, and 0.6% for MEK1 (**Figure S15**). Each of these also shows remarkable mutual exclusivity relative to the others in breast cancer patients reflecting oncogene functional redundancies. Further *in silico* comparison of oncogenic mutational occurrence across all cancer types (Cerami et al., 2012; Gao et al., 2013) indicate a strong mutual exclusivity between BRAF and KRAS, and a slight mutual exclusivity with EGFR (**Figure S16**).

### Alterations in transporter expression

One of the most pronounced oncogene-induced changes we observed is to proteins involved in solute transport that reverse upon MEK inhibition including: upregulation of SLC2A1, SLC6A15, SLC7A1, SLCO4A1, MF12/melanotransferrin, and down regulation of SLC22A5 and STEAP4 (**Figure 3D**). SLC6A15 acts as a preferential methionine amino acid transporter and shows dramatic upregulation in both KRAS and HER2 transformed cells. This is consistent with recent studies showing KRAS transformed cells exhibit an extreme sensitivity to methionine deprivation (De Sanctis et al., 2016). The Warburg Effect preferentially shifts cancer cells towards glycolysis, thereby promoting accelerated growth and division (Liberti and Locasale, 2016). We find SLC2A1, also known as glucose transporter-1 (GLUT1), is significantly upregulated in all oncogene transformed cells which would facilitate anaerobic glycolysis and increased growth for these cells. Our data show upregulation of SLCO4A1, a part of the organic anion transporter family that assists in transporting hormones such as prostaglandins, and vasopressin. Heightened expression of this hormone transporter has been previously seen in metastatic colorectal cancer (Buxhofer-Ausch et al., 2013). SLC7A1 has been shown to be upregulated across many cancer types, including colorectal cancer (Lu et al., 2013). Others have shown that SLC7A1 mRNA expression level decreases upon MAPK pathway inhibition(Calder et al., 2011), which is consistent with our proteomics data.

### General downregulation of receptor tyrosine phosphatases and tumor suppressors

Another important theme found within our data is a general down-regulation of surface proteins involved in receptor tyrosine phosphatases and other tumor suppressors such as: PTPRF, PTPRS, UNC5B, and BCAM (**Figure 2A, Figure 3D**). The expression of PTPRF and PTPRS, in particular, have been associated with decreased metastasis through the inactivation of EGFR signaling, emphasizing their importance as tumor suppressor proteins in numerous cancer contexts (Davis et al., 2018; Tian et al., 2018). Both of these proteins show a marked decrease in expression across the majority of the oncogene transformed cells, perhaps as a means to promote growth and metastasis. Similarly, UNC5B has recently been shown to halt tumor progression in an *in vivo* model of bladder cancer through inducing cell cycle arrest in the G_2_/M phase (Kong et al., 2016). Lastly, BCAM has been previously shown to be a tumor suppressor in a model of hepatocellular carcinoma (Akiyama et al., 2013), and decreased expression in all oncogene transformed cells may signify the removal of tumor suppressive signaling.

### Alteration in adhesion molecules

Metastasis is the leading cause of death for patients with cancer, and tumor cells acquire the ability to penetrate the surrounding tissues, thus leading to invasion (Martin et al., 2013). Many of these acquired functionalities are through changes in adhesion molecules on the cell surface, which play the role of mediating cell-cell interactions. Within our dataset, we identified five targets, LAMC2, LAMA3, LAMB3, PODXL, and MME, that are known to play important roles in mediating metastasis in various tumor types and we find to be upregulated across the majority of oncogenic mutants (**Figure 2A, Figure 3D**). LAMC2 has previously been shown to promote EMT and invasion in an *in vivo* model of lung adenocarcinoma (Moon et al., 2015). It has also been shown that colorectal cancer with high MAPK activity express heightened levels of LAMC2, supporting our proteomic results that show reversal upon MEK inhibition (Blaj et al., 2017). PODXL had increased expression in all six mutant cell lines, and has been implicated in increasing the aggressiveness of breast and prostate cancer through the induction of both MAPK and PI3K signaling (Sizemore et al., 2007). PODXL is also a glycoprotein with extensive mucin-type O-glycosylation, the overexpression of which can lead to increased metastatic potential through increased cell cycle progression via the PI3K and MAPK axes (Nielsen and McNagny, 2009; Paszek et al., 2014; Woods et al., 2017). MME, known as matrix metalloprotease-12, was found upregulated in all six oncogenic mutants compared to the empty vector control. MME has been reported to participate in the break-down of extracellular matrix. It is involved in tumor invasion in some settings (Kerkelä et al., 2002) and can be used to promote oncolytic viral infection in tumors (Lavilla-Alonso et al., 2012). Interestingly, MME is upregulated across all oncogenes in both the oncogene and MEKi datasets, suggesting that MME expression is not regulated through the MAPK or PI3K axis. This heightened expression of MME in all oncogenic contexts presents an opportunity for future research into synthetic lethality studies with MME inhibition in combination with MEKi or other MAPK targets.

### Immune Modulation

Another prominent cause of cancer progression is the evasion of immune surveillance. This can be achieved by overexpressing proteins that have a net immunosuppressive effect or by downregulating proteins that increase immune activation. Here, we identified three differentially regulated proteins, NT5E, HLA-F, and DSE, that play important roles in immune functions (**Figure 3D**). NT5E, which was upregulated in all MAPK mutants, promotes immunosuppression through the production of adenosine from AMP, which can decrease the capacity of natural killer cells to produce immune activating IFN molecules and prevents the clonal expansion of cytotoxic T cells in the surrounding tumor tissue (Gao et al., 2014). Conversely, HLA-F and DSE (also known as dermatan sulfate epimerase), were downregulated in all oncogene expressing cell lines, and both play a role in immune system activation and tumor-rejection, through activation of NK cells or cytotoxic T cells, respectively, in the tumor microenvironment (Dulberger et al., 2017; Liao et al., 2019). Both NT5E and HLA-F show reversed expression levels upon MEKi, suggesting a potential mechanism for immune re-activation after tumor development (**Figure 3D**).

### Down-regulation by shedding

Proteolytic cleavage of proteins at the cell surface leads to ectodomain shedding and the existence of neo N-termini. The loss of ectodomains would lead to decreased peptide detection by LC-MS/MS. We identified four surface proteins reduced in detection, MSLN, TACSTD2, NRCAM, and GPC1, that are known to undergo proteolysis to enhance the growth and metastasis of cancer. (Kawahara et al., 2017; Morello et al., 2016; Stoyanova et al., 2012). In particular, NRCAM, which had lower surface levels in all six oncogenic mutants is a cell-cell adhesion molecule; it is known to be shed by proteolysis, and the shed form stimulates cell proliferation via the AKT pathway (Conacci-Sorrell et al., 2005). Perhaps unsurprisingly, NRCAM did not exhibit a renormalization of expression levels after MEK inhibition, due to its exclusive function through the AKT pathway.

### Changes in glycosylation machinery and protein N-glycosylation

Alterations in glycosylation are common features of cancer cells (Pinho and Reis, 2015) which can result from a variety of factors, including changes in expression of glycosyltransferases, availability of sugar nucleotide substrates, alteration in expression of substrate proteins, or changes to the tertiary structure of the protein substrate that disrupt transfer of the glycan. We find glycosyl transferases FUT10, EXT2, GALNT11, GCTN2, and ST6GALNAC2 are highly upregulated across all oncogenic cell lines. FUT10 is an α-1,3-focusyltransferase, and FUTs are critical for the production of Lewis^X^ and Sialyl Lewis^X^ antigens which are hallmarks of invasion (Blanas et al., 2018). In fact, the most aggressive of our oncogene cell lines (HER2 and KRAS) are significantly upregulated in complex hybrid type glycans for which these are members, and especially noteworthy for ANPEP. EXT2 forms a heterodimer with EXT1 in the Golgi functioning as a glycosyl transferase involved in catalyzing the formation of heparin, a substrate that contributes to structural stability of the extracellular matrix, thereby mediating processes such as adhesion, immune infiltration, and signaling (Nagarajan et al., 2018). Unsurprisingly, alterations in heparan sulfate formation has been shown to have emerging roles in oncogenesis, of which EXT2 is a primary facilitator (Knelson et al., 2014). GALNT11 is a protein involved in the initiation of mucin-type O-glycosylation a well known marker that over-expressed in cancer (Villacis et al., 2016) ((Libisch et al., 2014) (Hussain et al., 2016). MUC-1 is also a very important drug target for immunotherapy (Bafna et al., 2010; Posey et al., 2016; Rivalland et al., 2015; Vassilaros et al., 2013). Interestingly, a common substrate for this family of enzymes, PODXL, is also upregulated across all oncogenic cell lines (see above), suggesting a possible mechanism by which increased GALNT11 promotes pro-growth phenotypes in these cell lines. GCTN2 is a glycosyltransferase (Bierhuizen et al., 1993), and specifically implicated in the transition from naïve to germinal center B cell (Meyer et al., 2018).

ST6GalNAc2 is a sialyltransferase that catalyzes attachment of GalNAc to core glycans and is paradoxically up-regulated in all the oncogene cell lines. A genome wide shRNA screen in mice for genes that suppress metastasis identified and validated ST6GalNAc2 as the strongest metastasis suppressor. ST6GalNAc2 is strongly down regulated in estrogen-receptor negative breast cancer tumors. Silencing of ST6GalNAc2 promotes binding of Galectin-3 and retention of tumor cells to sites of metastasis. Galectin-3 is a member of S-type lectins and known modulator of tumor progression (Liu and Rabinovich, 2005) by enhancing aggregation of tumor cells during metastasis and preventing anoikis (Bair et al., 2006; Grassadonia et al., 2002; Lin et al., 2015). Galectin-3 binds to core glycans branched glycan structures with terminal fucosylation on Galectin-3 binding protein (Gal-3BP) that ST6GalNAc blocks via sialylation (Lin et al., 2015). We found all glycopeptides from the eight glycosites Gal-3BP were down-regulated up to 50-fold in all cell lines (**Figure S14**).

Our glycoproteomic analyses show protein N-glycosylation is dramatically changed upon oncogenic transformation, and that distinct genetic drivers of oncogenesis promote unique changes to the glycoproteome. The majority of differentially expressed glycopeptides were unique to individual cell lines. We observed greatest similarity between BRAF^V600E^, EGFR^L858R^, and MEK^DD^, which has been consistent throughout different analyses of these cell lines (**Figure 5B**). There were significant differences between the oncogene Cluster 1 (KRAS/HER2) and Cluster 2 (EGFR/ BRAF/ MEK/AKT). Specifically, we found differential increase in complex hybrid glycans and decrease in high mannose in Cluster 1 and just the opposite for Cluster 2.

There is abundant evidence to suggest that high-mannose type glycans are present at the cell surface, and furthermore, that cancerous cells display increased abundance of high-mannose glycans at their cell surface (Holst et al., 2016; Hua et al., 2014). A similar observation was reported in the glycomic comparison of transformed versus human embryonic stem cells (hESC), where high-mannose glycans were observed at significantly higher abundance on plasma membranes of hESCs (An et al., 2012). Together, these observations help to explain the similarities between cancerous cells and hESCs, such as the regulation of tumor suppressor genes and the ability to self-renew (Shackleton, 2010). This is consistent with our studies although we cannot exclusively rule out that some of the high mannose glycans we observe are coming from proteins in route to the membrane.

Our data suggest glycosylation is quite heterogeneous possibly reflecting incomplete maturation. The most stunning example in our data of heterogeneity in glycosylation was for ANPEP, an amino peptidase also upregulated at the protein level across most cell lines. The most extreme example was site N234 of ANPEP where we found 29 different glycan structures. When these are arranged in order of glycan maturation, we observed a close representation of its steady state progression through the secretion pathway (**Supplementary Figure 13**). ANPEP also showed the greatest representation within complex/hybrid glycopeptides that were upregulated in HER2 and KRAS^G12V^ cell lines (**Figure 5D**). ANPEP was most upregulated in KRAS^G12V^ in surfaceomic analyses and its expression was not dramatically influenced by MEK inhibition. ANPEP expression in our surfaceomic and glycoproteomic data represents one way in which KRAS^G12V^ diverges from other members of the MAPK pathway. These data, combined with ANPEP’s previously identified role in angiogenesis(Pasqualini et al., 2000), underscore its importance in KRAS^G12V^-positive tumors. ANPEP glycosylation may serve as a very sensitive biomarker of glycan metabolism in cells given its highly diverse and heterogeneous patterns.

## Conclusions

We provide a large-scale comparative study of how six neighboring proliferative oncogenes cause large cell autologous remodeling in the surface glycoproteome. While many of the changes are specific to given oncogenes at both the protein and glycopeptide level, we observe common biological themes suggesting functional redundancy of the specific cellular expression. This is consistent with observations that common tumors can express these different oncogenes although not together. We commonly observe upregulation of surface proteins involved in metabolite transport, glycosylation, and immune suppressors and down regulation of adhesion proteins and tumor suppressors which is consistent with increased cell growth and invasion that are well-known properties of cancer cells. These studies were deliberately conducted in isogenic cell lines to isolate the oncogene-induced changes. Even in a simplified autologous cell model we recapitulate many of the hallmarks of transformed cells driven by complex but functionally redundant changes to the surfaceome. We believe the work provides important insights into the similarities and differences among the neighboring oncogenes and provide new opportunities to pursue antibody tools to follow in more complex tumor settings.

## Acknowledgements

We wish to thank Professor Sourav Bandyopadhyay at UCSF for some of the lenti constructs and cell lines, Michael Westphall, Evgenia Shishkova, and Jean Lodge at University of Wisconsin for instrumentation and helpful discussions. JAW is grateful to funding from the National Institutes of Health grant R35GM122451 and Celgene corporation. KKL thanks Canadian Institutes of Health Research (CIHR) for post-doctoral funding. LLK is indebted to the Chemistry and Chemical Biology Training grant (T32 GM064337) at UCSF for funding. JJC wishes to thank NIH for generous funding from P41 (GM108538, The National Center for Quantitative Biology of Complex Systems). GMW was supported in part by the Biotechnology Training Program (5 T32 GM008349). NMR gratefully acknowledges support from National Institutes of Health grant K00CA21245403.

## Materials and Methods

All MCF10A cells were cultured according to established protocols unless otherwise stated. Surfaceome of each oncogene transformed cell line was compared to empty vector (EV) control using SILAC based quantification. For inhibition study, each oncogene transformed cell line was treated with 100nM MEK inhibitor PD032590 or DMSO control for three days and surfaceome changes were compared in each respective cell line using SILAC based quantification. Glycoproteome of each oncogene transformed cell line was quantified using label-free quantification. Detailed materials and methods are included in SI appendix.

