## Supplemental Text and Figures for "Broad and thematic remodeling of the surface glycoproteome on isogenic cells transformed with driving proliferative oncogenes"

**This PDF file includes:**

Supplemental Materials and Methods
Figures S1 to S16
SI References

### Supplemental Materials and Methods

**Cell line generation**

Parental MCF10A, MCF10A$^{EV}$ (empty vector with puromycin resistant marker), MCF10A Her2, MCF10A KRAS$^{G12V}$, MCF10A BRAF$^{V600E}$, MCF10A myristoylated AKT (AKT$^{myr}$) were a gift from Sourav Bandyopadhyay (1). The lentiviral transfer plasmids pLX304 EGFR$^{L858R}$ and pLX302 MEK$^{S218D/S222D}$ (MEK$^{DD}$) were a gift from Sourav Bandyopadhyay (1). MCF10A EGFR$^{L858R}$ and MEK$^{DD}$ cell lines were generated by lentiviral transduction of the aforementioned plasmids into parental MCF10A cell lines. After transduction, cells were allowed to recover for 2 days prior to puromycin resistance selection for 1 week. Cells were grown in DMEM supplemented with 5% horse serum (Invitrogen #16050-122), 20 ng/ml EGF, 0.5 mg/ml hydrocortisone, 100 ng/ml cholera toxin, and 10 µg/ml insulin.

**Growth inhibition by MEK inhibitor**

For cellular growth studies, each cell line was seeded at 5000 cells per well in a white 96-well plate with and without 100 nM MEK inhibitor (PD032590, Selleck Chemicals), and with and without growth factor (hEGF and insulin) in triplicate. Each day, one plate was assayed using CellTiter-Glo (Promega) according to standard protocol. Luminescence was measured using a Tecan plate reader Infinite 200 PRO and data was analyzed using GraphPad Prism 8 (La Jolla California USA, www.graphpad.com)

**SILAC labeling and Surfaceome analysis**

For surfaceome analysis quantified using SILAC, all cell lines were grown were cultured in DMEM SILAC media (Thermo Fisher Scientific; Waltham, MA) containing L-[$^{13}C_6$,$^{15}N_2$]lysine and L-[$^{13}C_6$,$^{15}N_4$] arginine (heavy label; Cambridge isotope laboratories, Tewksbury, MA) or L-[$^{12}C_6$,$^{14}N_2$]lysine and L-[$^{12}C_6$,$^{14}N_4$]arginine (light label) for 5 passages to ensure full incorporation of the isotope labeling on cells.

In order to compare oncogene transformed cells with viable empty vector cells, all cells were grown in the presence of growth factors. After seven passages in SILAC media, $20x10^6$ cells were harvested at 80% confluence, mixed at 1:1 cell count ratio, and subjected to the CSC protocol as modified to include all tryptic fragments. All experiments were performed in duplicate in both forward and reverse SILAC labeling scheme such that a total of four biological replicates were analyzed. Similarly, to understand the effect of the MAPK signaling pathway, each cell line was treated with 100nM MEK inhibitor (PD032590) or 1% DMSO for three days. Approximately $10 \times 10^6$ cells were harvested from each condition and mixed at a 1:1 ratio for CSC enrichment. Inhibitor experiments were performed with forward and reverse SILAC labeling such that a total of two biological replicates were analyzed.

To enrich for the surfaceome, live cells were treated with a sodium periodate buffer (2 mM NaPO4, PBS pH 6.5) at 4°C for 20 mins to oxidize glycoproteins. Aldehydes generated by periodate oxidation were then reacted with biocytin hydrazide in a labeling buffer (1 mM biocytin hydrazide (biotium), 10 mM analine (Sigma), PBS pH 6.5) at 4°C for 90 mins. Cells were then washed four times in PBS pH 6.5 to remove excess biocytin-hydrazide and flash frozen. Frozen cell pellets were lysed using RIPA buffer (VWR) with protease inhibitor cocktail (Sigma-Aldrich; St. Louis, MO) at 4°C for 30 mins. Cell lysate was then sonicated, clarified, and incubated with 500µL of neutravidin agarose slurry (Thermo Fisher Scientific) at 4°C for 30 mins. The neutravidin beads were then extensively washed with RIPA buffer, high salt buffer (1M NaCl, PBS pH 7.5), and urea buffer (2M urea, 50mM ammonium bicarbonate) to remove non-specific proteins. Samples were then reduced on-bead with 5mM TCEP at 55°C for 30 mins and alkylated with 10mM iodoacetiamide at room temperature for 30 mins. To release bound proteins, we first performed an on-bead digestion using 20µg trypsin (Promega; Madison, WI) at room temperature overnight. The "tryptic" fraction was then eluted using spin column and the neutravidin beads were extensively washed again with RIPA buffer, high salt buffer (1M NaCl, PBS pH 7.5), and urea buffer (2M urea, 50mM ammonium bicarbonate). To release the remaining trypsin digested N-glycosylated peptides bound to the neutravidin beads, we performed a second on-bead digestion using 2500U PNGase F (New England Biolabs; Ipswich, MA) at 37°C for 3 hrs. Similarly, the "PNGase F" fraction was eluted using a spin column. Both tryptic and PNGase F fractions were then desalted using SOLA HRP SPE column (Thermo Fisher Scientific) using standard protocol, dried, and dissolved in 0.1% formic acid, 2% acetonitrile prior to LC-MS/MS analysis.

**SILAC surfaceome mass spectrometry analysis**

Approximately 1µg of peptide was injected to a pre-packed 0.75mm x 150mm Acclaimed Pepmap C18 reversed phase column (2µm pore size, Thermo Fisher Scientific) attached to a Q Exactive Plus (Thermo Fisher Scientific) mass spectrometer. For "tryptic" fraction, peptides were separated using a linear gradient of 3-35% solvent B (Solvent A: 0.1% formic acid, solvent B: 80% acetonitrile, 0.1% formic acid) over 180 mins at 300µL/min. Similarly, the "PNGase F" fraction was separated using the same gradient over 120 mins. Data were collected in data-dependent mode using a top 20 method with dynamic exclusion of 35 secs and a charge exclusion setting that only sample peptides with a charge of 2, 3, or 4. Survey scans were collected as profile data with a resolution of 140,000 (at 200 m/z), AGC target of 3E6, maximum injection time of 120 ms, and scan range of 400 - 1800 m/z. MS2 scans were collected as centroid data with a resolution of 17,500 (at 200 m/z), AGC target of 5E4, maximum injection time of 60 ms with normalized collision energy at 27, and an isolation window of 1.5 m/z with an isolation offset of 0.5 m/z.

**SILAC surfaceome data processing**

Peptide search for each individual dataset was performed using ProteinProspector (v5.13.2) against 20203 human proteins (Swiss-prot database, obtained March 5, 2015) with a false discovery rate (FDR) of <1%. To estimate the efficiency of the surface proteome enrichment method, a list of extracellular proteins was generated by searching for "membrane" but not "mitochondrial" or "nuclear" using uniprot subcellular localization annotations. We found ~60% of peptides identified in the tryptic fraction and ~90% of peptides identified in the PNGase F fraction belonged to the extracellular proteome reflecting a high and expected enrichment ratio. All protein identifications were then filtered using the same list to ensure a stringent assignment of unique peptides.

Quantitative data analysis was performed using Skyline (2) software using the MS1 filtering function. Specifically, spectral libraries from forward and reverse SILAC experiments were analyzed together such that MS1 features without an explicit peptide ID would be quantified based on aligned peptide retention time. The first four isotopic peaks of precursor ions were then quantified at 50% FWHM defined by ms1 scanning resolution of 140,000 (at 200 m/z). The boundary of peptide elution time was determined by default algorithm and the total peak area was used as the peptide quantification value. An isotope dot product of at least 0.8 (as calculated by Skyline) was used to filter out low quality peptide quantification, and a custom report was generated for further processing and analysis using R. To ensure stringent quantification of the surface proteome, several filters were applied to eliminate low confidence protein identifications. In the tryptic fraction, only peptides with five or more well quantified peptides were included. In the PNGase F fraction, only peptides N to D deamidation modification were included. Forward and reverse SILAC datasets were then combined and reported as median log2 enrichment values. All data analyses were carried out using R: A language and environment for statistical computing (3). Specifically, gene set enrichment analysis was carried out using fast pre-ranked gene set enrichment analysis (fgsea) package from Bioconductor (4).

**Glycoproteome mass spectrometry sample preparation**

Cell lines were analyzed in biological triplicate. $20 \times 10^6$ cells were suspended in 200 µL 6 M guanidine HCl and boiled for 5 min at 100 °C. Protein was precipitated by the addition of 1,800 µL methanol and pelleted by centrifugation at 12,000 x G for 5 min. Pelleted protein was resuspended in lysis buffer (8M urea, 40 mM 2-chloroacetamide, 10 mM tris(2-carboxyethyl)phosphine, 100 mM tris pH 8) and incubated for 10 min at RT before diluting to [urea] < 2M with 50 mM tris. Trypsin was added at a protein:enzyme ratio of 100:1 and incubated overnight at RT with gentle rocking. After digesting overnight, the solution was adjusted to pH < 2 and desalted with StrataX reverse phase SPE cartridge (Phenomenex, Torrence, CA). Eluted peptides were dried under reduced pressure and quantified by bicinchoninic acid assay (Pierce™ Quantitative Colorometric Peptide Assay, Thermo Fisher Scientific, Waltham, MA).

**Lectin Glycopeptide enrichment**

To enrich for glycopeptide, 200 µL of ConA (Agarose bound ConA, Vecetor Labs, Burlingame, CA) was washed 3X with 500 µL wash buffer (50 mM Tris, pH 8) and combined with 2 mg tryptic peptides in 500 µL wash buffer. Solution was rocked end-over-end for 1.5 hr at 4 °C. Agarose beads were washed 8X with wash buffer. Glycopeptides were eluted 3X with 500 µL ConA elution buffer (200 mM α-methylmannoside, 1% trifluoroacetic acid). 500 µL of elution was portioned for deglycosylation with protein N-glycosidase (New England Biolabs, Ipswich, MA). The remaining eluent was desalted with StrataX reverse phase cartridges (Phenomenex, Torrence, CA). The deglycosylated sample was desalted after overnight incubation with PNGase (New England Biolabs, Ipswich, MA) at 37 °C.

**HILIC glycopeptide enrichment**

A method adapted from Totten et al., 2017 (but with different exchange resin) was used. A mixed mode strong anion exchange (Oasis MAX, 60 mg, Waters Corporation, Milford, MA) was equilibrated with 1) 1 mL acetonitrile 2) 1 mL 100 mM triethylammonium acetate 3) 3 mL water, and 4) 1 mL 95% acetonitrile, 1% trifluoroacetic acid. 3.5 mg of tryptic plasma peptides was loaded in 200 uL 50% acetonitrile, 0.1% trifluoroacetic acid diluted in 3 mL 95% acetonitrile, 1% trifluoroacetic acid. The column was washed with 3 mL 95% acetonitrile, 1% trifluoroacetic acid before eluting into a clean vial with 3x1 mL 50% acetonitrile, 0.1% trifluoroacetic acid. 500 µL of elution was portioned for deglycosylation as above.

Our data confirm that the ConA and HILIC enrichments allowed for sampling of different glycan classes and when combined allowed for greater number of identifications (**Figure S9**). As expected for ConA, almost 80% of the identifications contained high mannose or paucimannose type glycans, whereas these high mannose classes comprised only 45% of the identifications in the HILIC dataset (**Figure S9A**). 1,635 unique glycopeptide identifications were made from ConA enrichments and 1,544 were identified from the HILIC enrichments, yet only 541 of these overlapped (**Figure S9B**). Similarly, 518 glycosites were characterized in the ConA dataset, while 545 glycosites were identified in the HILIC dataset, with 278 glycosites shared between them (**Figure S9C**). Glycopeptide mass, charge, glycan and peptide backbone mass, amino acids per peptide and hydrophobicity were all generally lower for ConA enrichments compared to HILIC (**Figure S9D-I**). These data are consistent with the lectin specificity of ConA for mannose and the reduction of glycan-type bias of the HILIC enrichment. The two enrichment methods are indeed complementary and together provide greater glycopeptide coverage.

**Glycoproteome mass spectrometry analysis**

All samples were analyzed on an ETD-enabled Q-IT-OT tribrid mass spectrometer (Orbitrap Fusion Lumos, Thermo Fisher Scientific, San Jose, CA) that had previously been modified to perform activated ion electron transfer dissociation (AI-ETD) (Riley et al., 2017) and was operated under data dependent acquisition following nano-LC separation and electrospray ionization. Instrument parameters were changed depending on sample type. For the analysis of unenriched and deglycosylated samples, MS1 survey scans were performed in the Orbitrap (240K resolving power, scan range 300-1,350 m/z, maximum injection time 50 ms, AGC target 1e6). Peptide precursors were isolated with the quadrupole (isolation width 0.7 m/z), subjected to HCD fragmentation (NCE 25%), and resulting fragment ions were analyzed by the ion trap (Rapid scan rate, scan range 200-1,200 m/z, maximum injection time 18 ms, AGC target 1e4). A dynamic exclusion duration of 20 s was enabled. Survey scans of intact glycopeptide enriched samples were performed in the Orbitrap (60K resolving power, scan range 600-1,800 m/z, maximum injection time

50 ms, AGC target 5e5). Peptide precursors were isolated with the quadrupole (isolation width 1.8 m/z), subjected to HCD fragmentation (NCE 30%), and resulting fragment ions were analyzed by the Orbitrap (30K resolving power, scan range 120-1,200, maximum injection time 60 ms, AGC target 5e4). HCD scans containing an oxonium ion within the top 20 most intense ions triggered an MS2 scan of the same precursor with AI-ETD fragmentation. Product ions were analyzed by the Orbitrap (30K resolving power, scan range 150-3,000 m/z, maximum injection time 400 ms, AGC target 3e5, calibrated reaction time). A dynamic exclusion duration of 60 s was enabled.

**Glycoproteome database searching**

Raw data from the analyses of unenriched and deglycosylated samples were searched with the MaxQuant Quantitative Software Suite against a database of human proteins downloaded from Uniprot on February 7, 2017.(5) Methionine oxidation and asparagine deamidation were allowed as variable modifications. Carbamidomethylation of cysteines was imposed as a fixed modification. Label free quantitation and intensity based absolute quantitation were performed.(6, 7) Raw data from intact glycopeptide analyses were searched with the Byonic™ search engine (Protein Metrics, Cupertino, CA) against a database of proteins for which deglycosylated peptides were identified at a N-glycosylation motif. A database of 182 N-glycans was provided as variable modification of asparagine with N-glycosylation sequons. Data were filtered, annotated, and organized with in-house software. Specifically, PSMs with Byonic scores below 150, delta mod scores below 10, log probabilities below 1, peptide length below 6, or multiply glycosylated were removed (**Figure S8**).

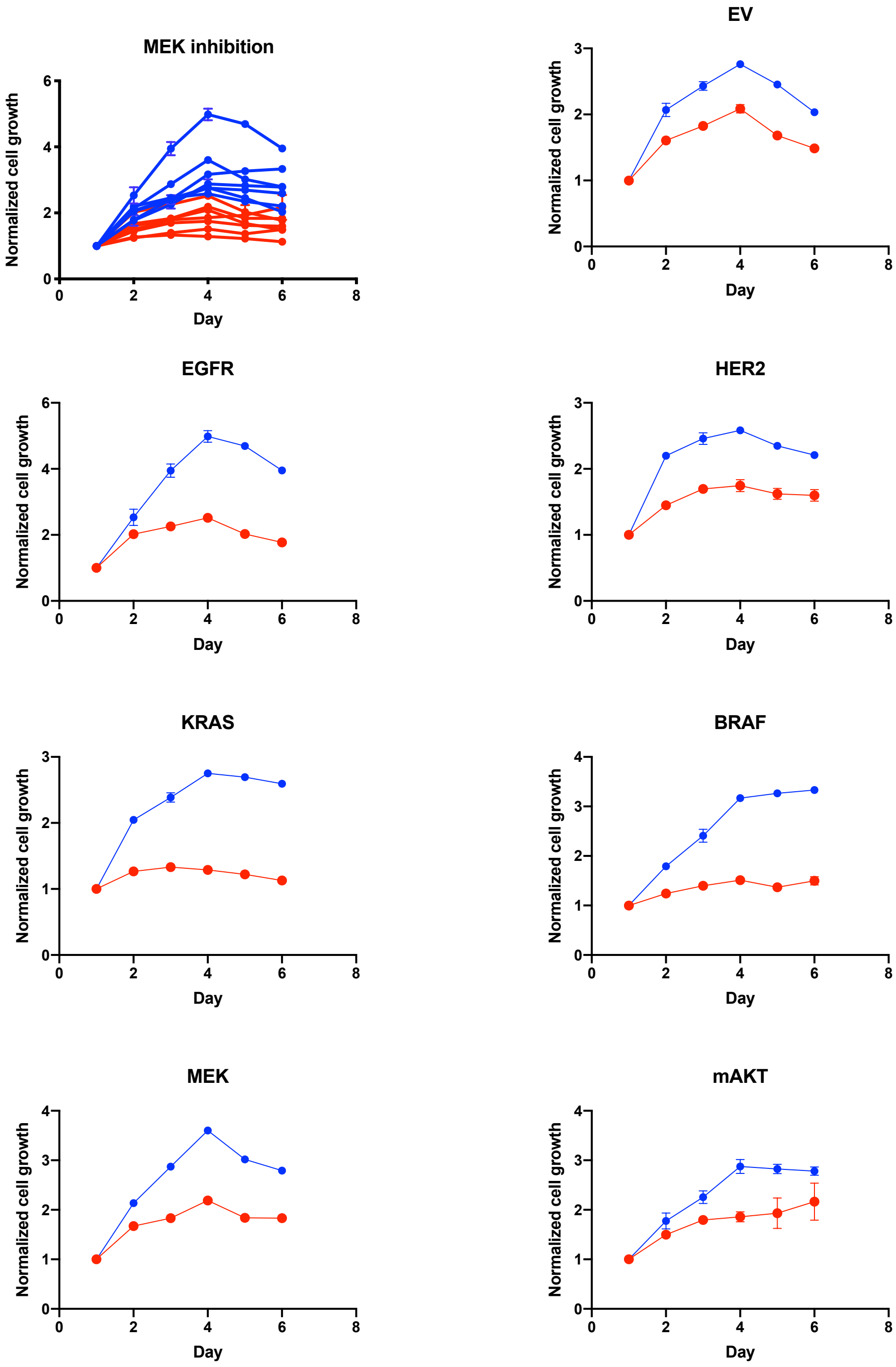


**Figure S1:** Growth inhibition by MEK inhibitor. All normalized growth curves are plotted together in the first panel treated with 100nM MEK inhibitor (PD0325901) in red and DMSO treatment control in blue. All following panels show each cell line individually. Data show average values of 3 replicates, and error bars show one standard deviation.

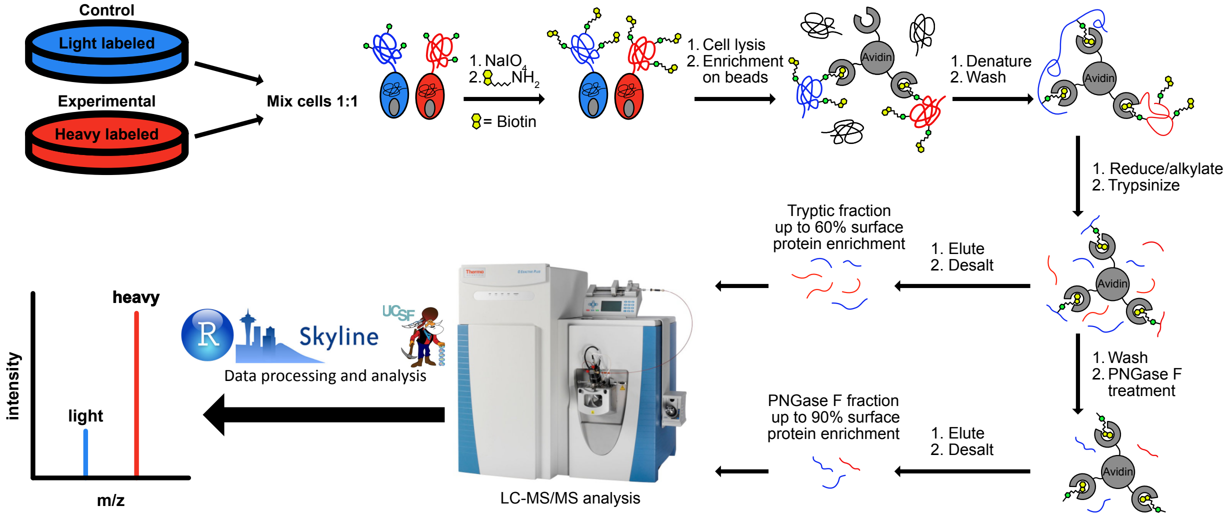


**Figure S2:** Cell surface capture sample preparation schematic and peptide quantification using SILAC. Heavy or light labeled cells were mixed at a 1:1 ratio prior to biotinylation of glycans using hydrazide chemistry. Proteins enriched on neutravidin beads were washed extensively and trypsinized on beads followed by PNGase treatment yielding two fractions. Peptides were analyzed using a Q Exactive Plus instrument followed by data analysis using Protein Prospector, MS1-level quantification using area under the curve integration in Skyline, and R. See methods section for detailed description.

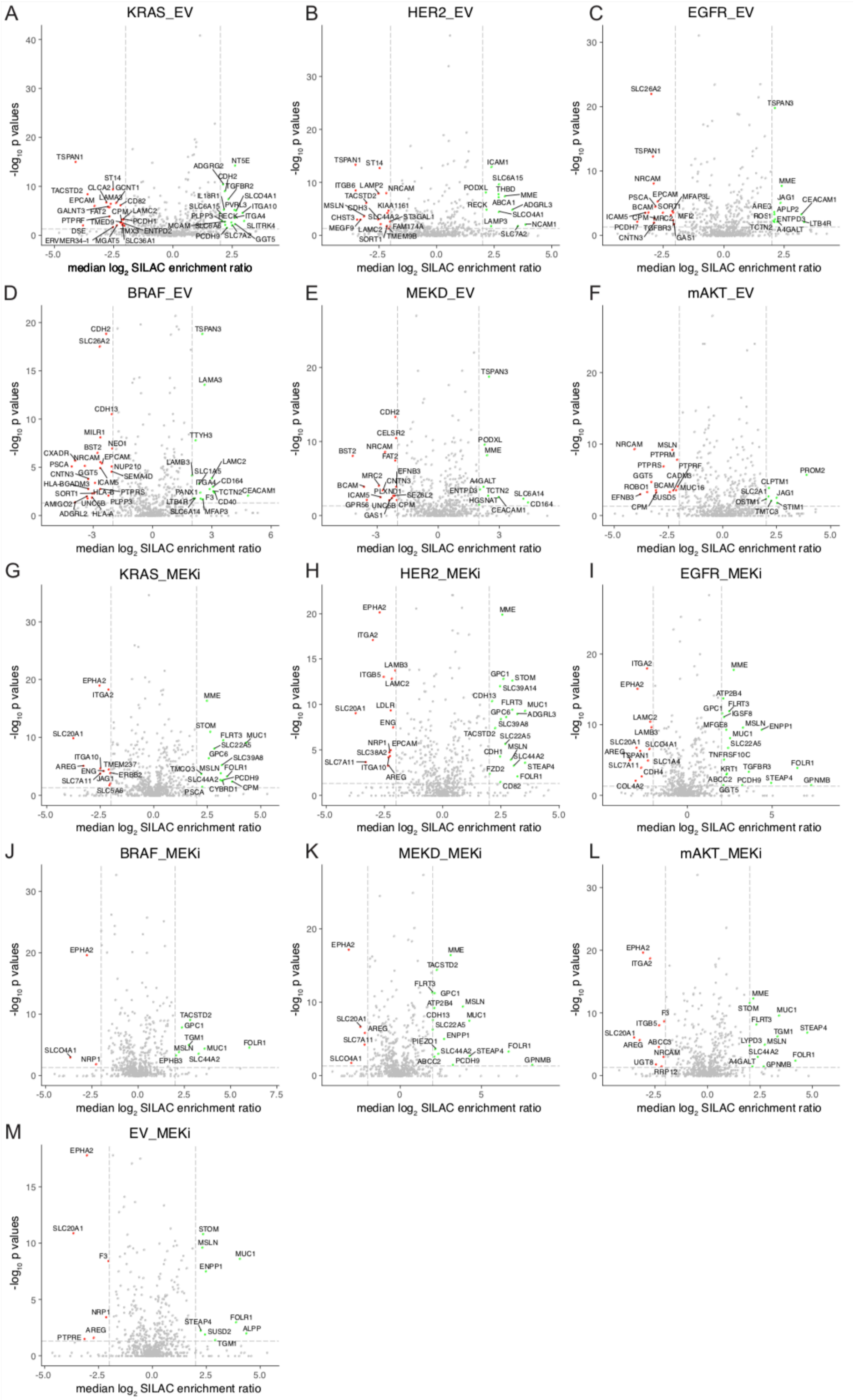

**Figure S3:** Volcano plots of each experiment. (A-F) Volcano plots of oncogene transformed MCF10A versus empty vector control. (G-M) Volcano plots of oncogene transformed MCF10A cultured with 100nM MEK inhibitor (PD0325901) treatment or 1% DMSO for 3 days. Dotted lines show fold change > 4 and p < 0.05.

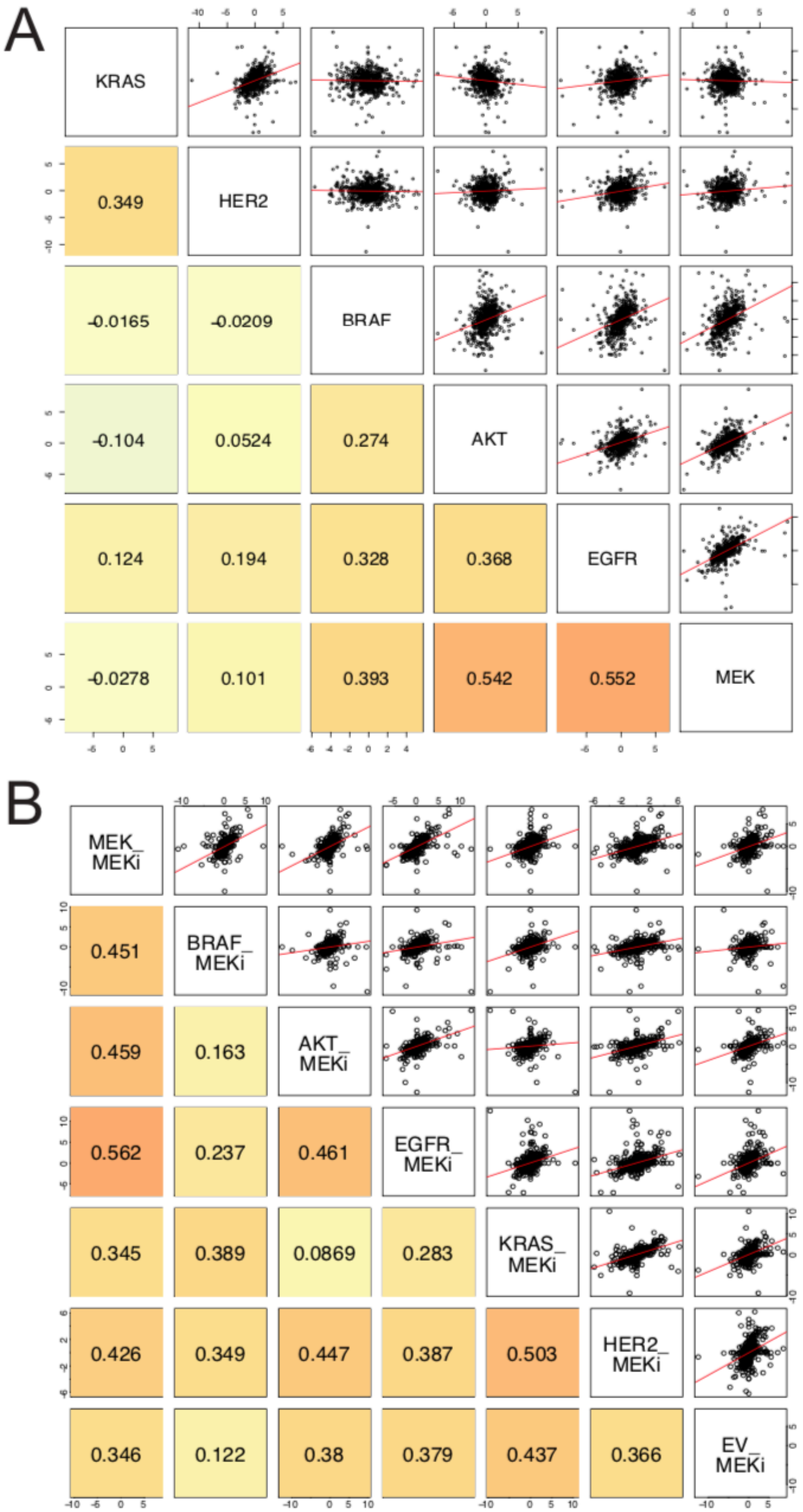


**Figure S4**: (A) Pairwise comparison of SILAC enrichment ratio between all oncogenic cell lines vs empty vector control reveals two cluster of correlation. (B) Pairwise comparison of SILAC enrichment ratio between all cell lines treated with MEK inhibitors showed high overall correlation.

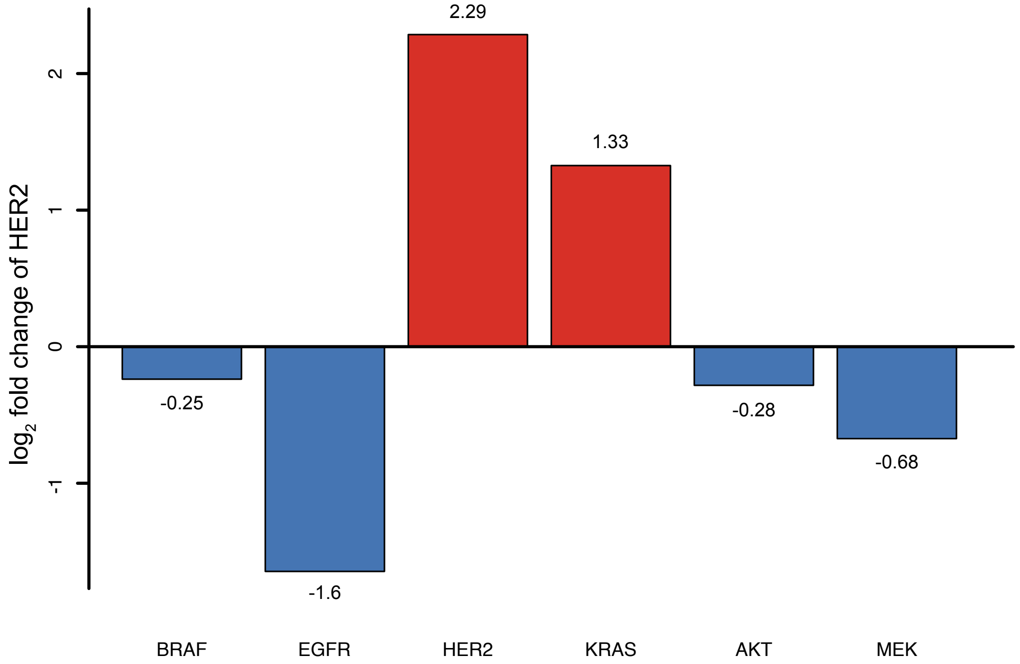


**Figure S5:** HER2 is up-regulated in MCF10A transformed with KRAS$^{G12V}$, and down-regulated in MCF10A transformed with EGFR$^{L858R}$. $Log_2$ fold change is indicated by number on top or below the bar.

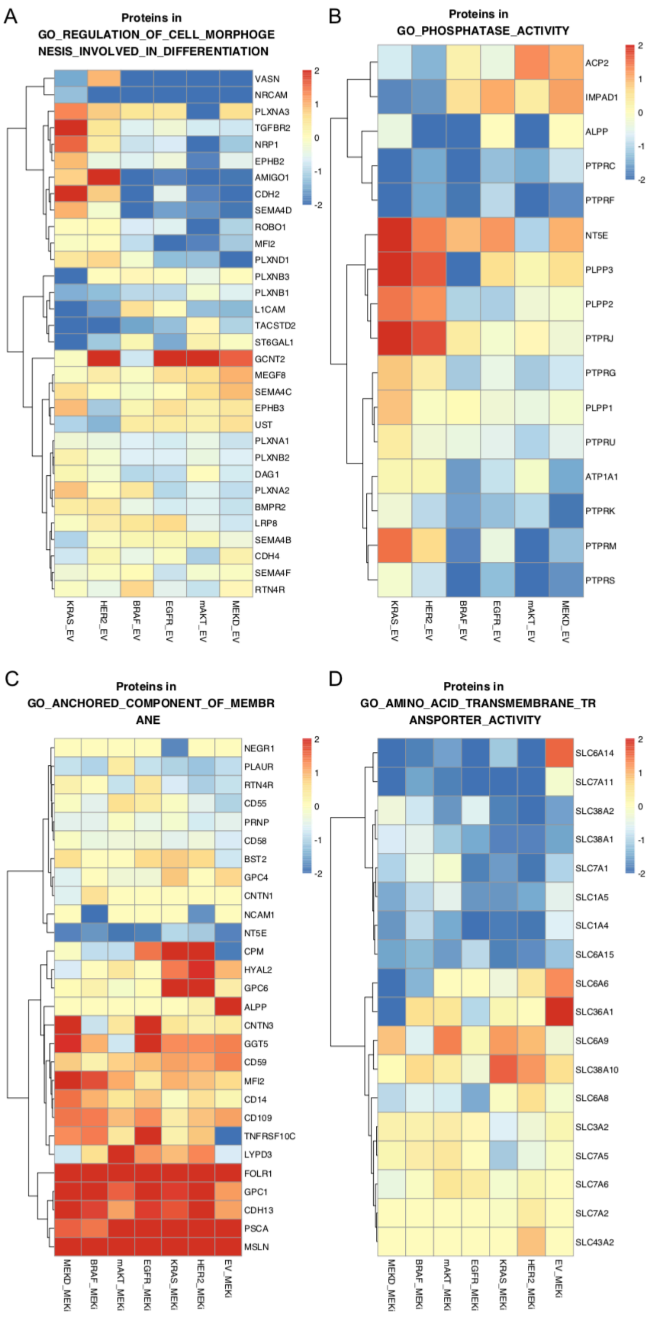


**Figure S6:** Selected gene sets are highlighted to illustrate differentially regulated proteins that contributes to different clusters of transformed MCF10A (A-B) or under MEK inhibition (C-D). (A) Proteins involved in regulation of cell morphogenesis involved in differentiation (GO: 0010769) are down regulated all cells. (B) Proteins with phosphatase activities (GO: 0016791) are up-regulated in the KRAS and HER2 cluster (blue) while the same proteins are down-regulated in the other oncogenes cluster. PTPRC and PTPRF in particular is down-regulated in all cells. (C) Cell anchored component of membranes (GO: 0031225) is up-regulated upon MEK inhibition. (D) Amino acid transmembrane transporter activity (GO: 0015171) is down-regulated up on MEK inhibition.

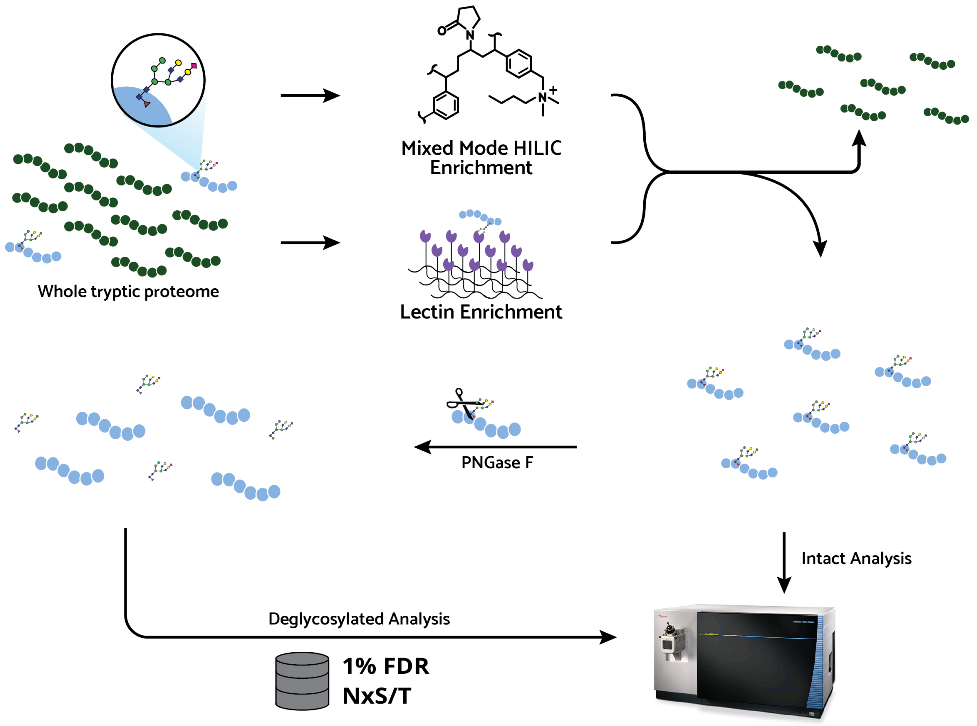


**Figure S7:** A workflow for intact and deglycosylated glycopeptide analysis is depicted. Two mg of tryptic proteomes are enriched with 1) a mixed-mode strong anion exchange cartridge with HILIC mobile phases (mm-SAX-HILIC) and 2) agarose-bound Concanavalin A. One quarter of each enriched yield is separated for deglycosylation with PNGase F and analyzed by LC-MS/MS with HCD fragmentation. Intact samples are analyzed directly on an AI-ETD-enabled LC-MS/MS with oxonium ion-triggered AI-ETD fragmentation.

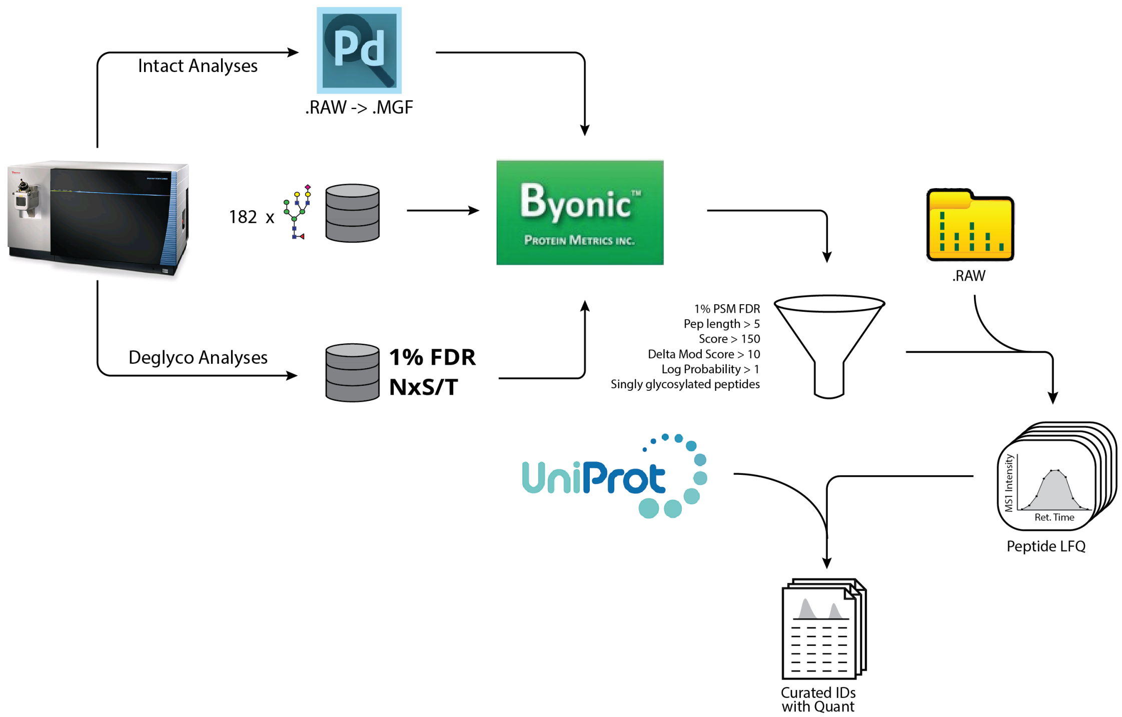


**Figure S8:** A workflow for intact glycopeptide identification is depicted. Raw data are converted to a generic text format using a conversion node within Proteome Discoverer 2.1. Tandem mass spectra are searched with the Byonic search engine against a database of forward and reverse sequences of proteins identified as having a deamidated asparagine at an N-glycosylation motif (NxS/T) in deglycosylated analyses. A library of 182 N-glycan compositions is provided as possible variable modifications. Search results are further filtered to ensure only confident assignments remain. Quantitation is provided for each glycopeptide by integrating the area under the curve of the eluting peptides from MS1 scans. Identifications are further annotated with pertinent information from the Uniprot database.

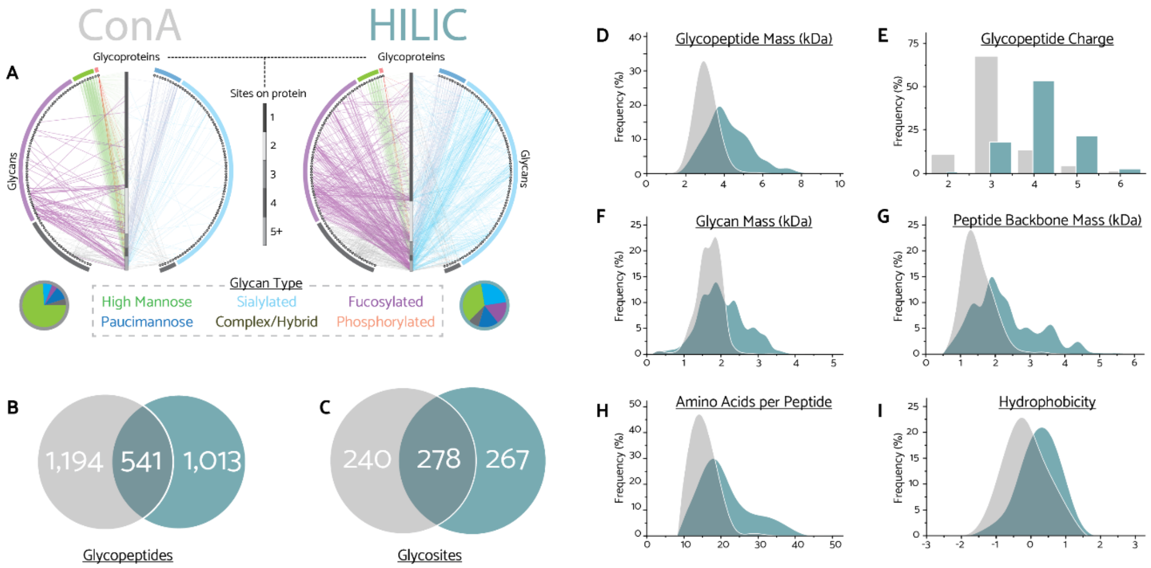


**Figure S9:** Comparing the glycoproteome subpopulations that are captured from Concanavalin A (gray) and mixed mode strong anion exchange hydrophilic interaction (mm-SAX-HILIC) (blue) glycopeptide enrichment. **a)** Glycan-glycoprotein network map. Glycan nodes are organized around the perimeter, sorted by type, while identified glycoproteins are arranged on the vertical axis and organized by the number of observed glycosites it harbored. An edge is drawn between a glycan and its modified glycoprotein for observed glycopeptides. **b)** Overlap of unique glycopeptides identified from each enrichment. **c)** Unique glycosites from each enrichment method. Distributions of **d)** glycopeptide mass, **e)** observed charge states, **f)** glycan mass, **g)** peptide backbone mass, **h)** peptide length, and **i)** Kyte-Doolittle hydrophobicity are displayed for each enrichment.

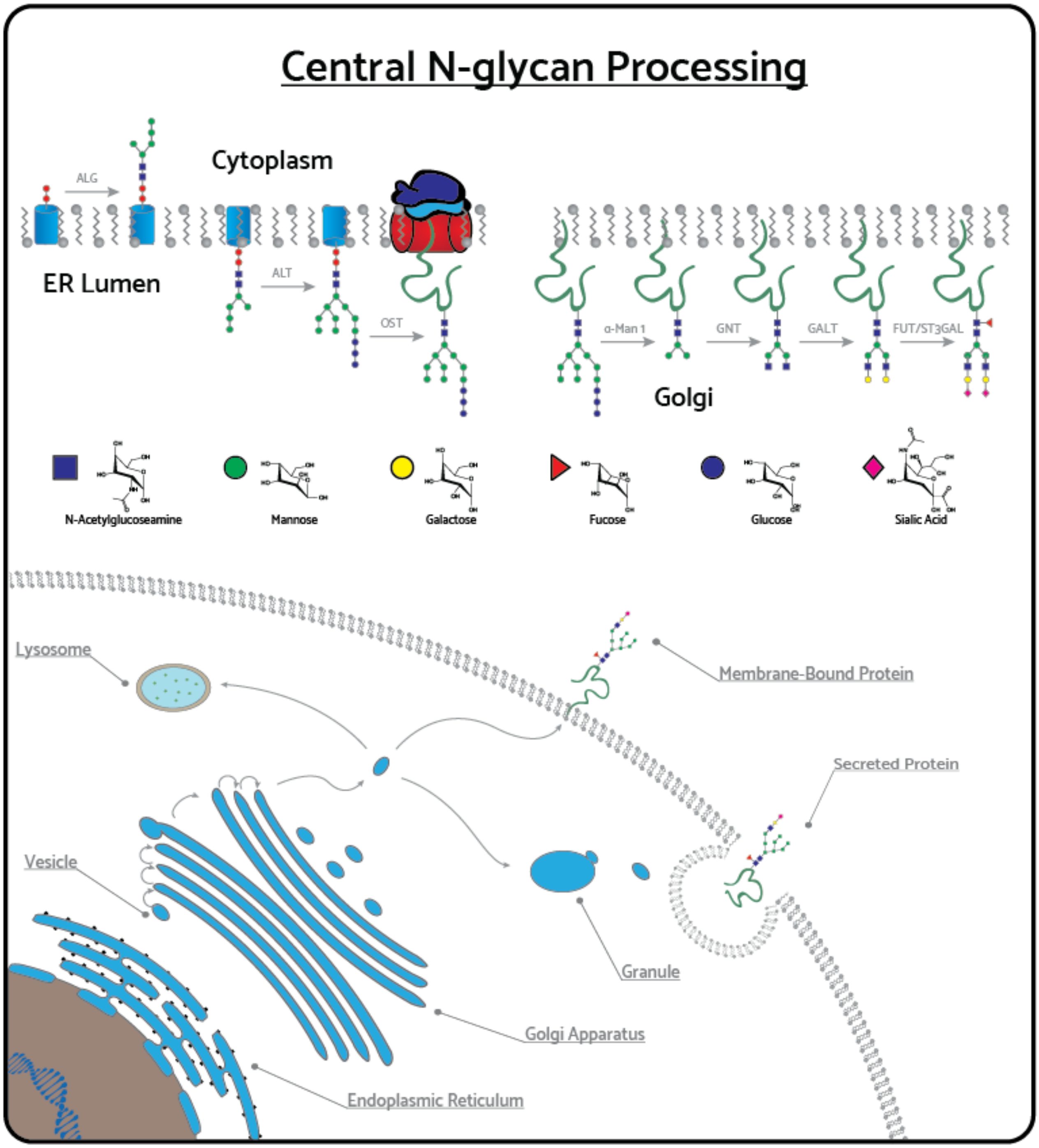


**Figure S10:** A representation of central N-glycan maturation through the endoplasmic reticulum, Golgi apparatus, and secretion pathway.

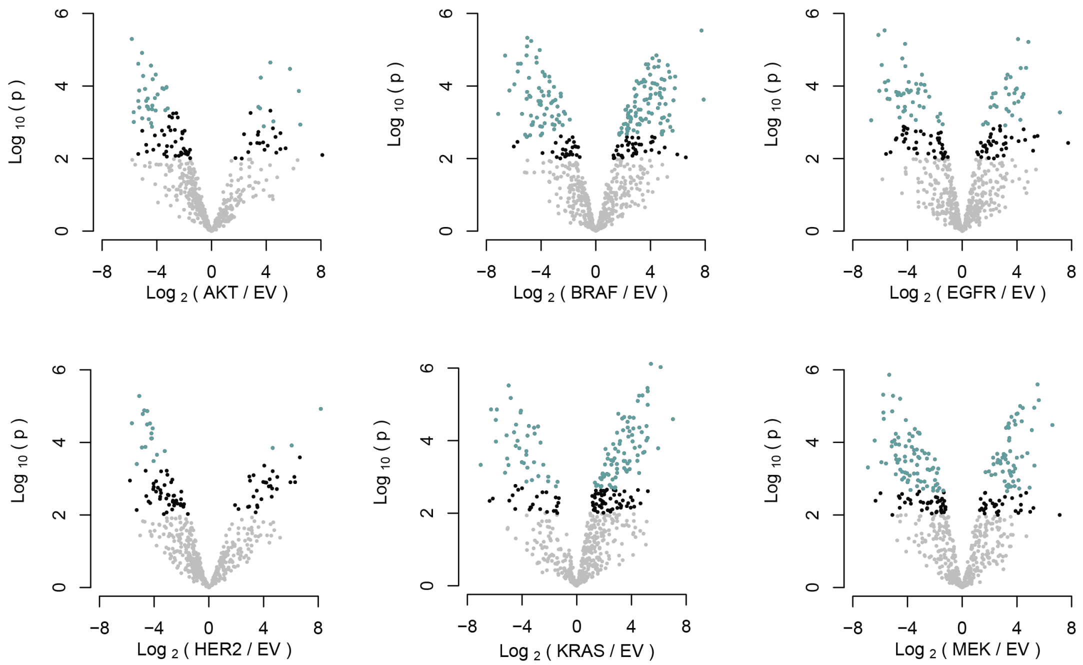


**Figure S11:** Differential expression of glycopeptides observed in each cell line compared to empty vector control. Significant values ($p < 0.01$) are in black. Those that pass below a 5% false discovery rate by Bonferroni correction are shown in green.

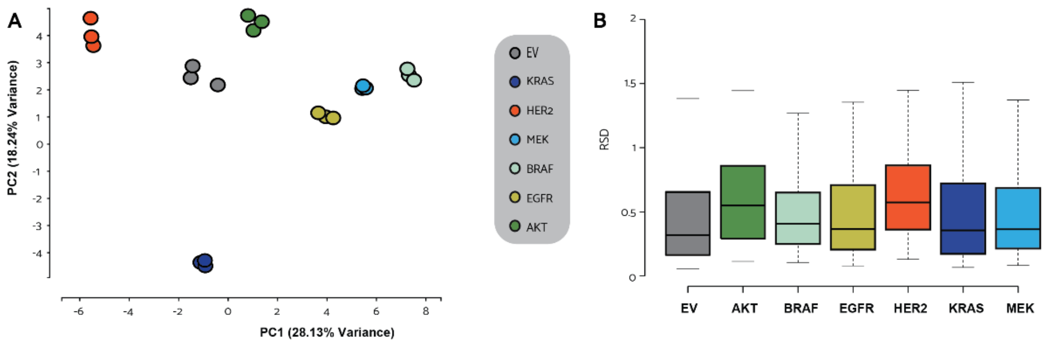


**Figure S12:** Figures of merit for quantitative glycopeptide analyses. **a)** Separation of individual cell lines and clustering of biological replicates is represented by a scatter plot of the first two principal components. **b)** Box plots show distributions of relative standard deviations across all glycopeptide measurements within each cell line.

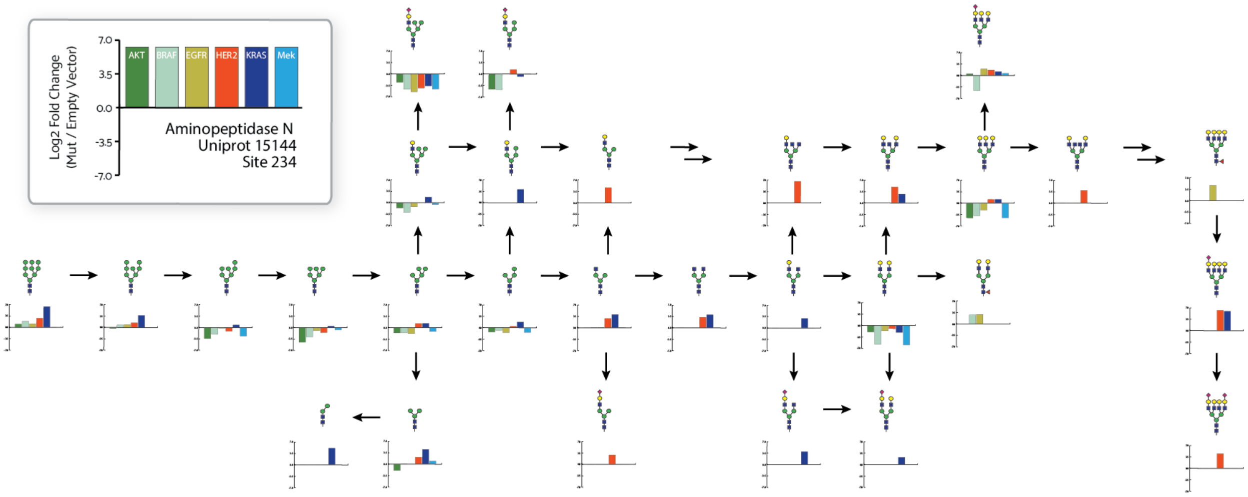


**Figure S13:** Glycopeptides identified from site N234 of aminopetidase N, on which the highest degree of microheterogeneity was observed. Glycans are arranged in the order of maturation, starting from the least mature, Hex(9)HexNAc(2). The fold change compared to empty vector control is provided in a bar plot below each glycan illustration. The legend in the upper left provides the color associated with each cell line.

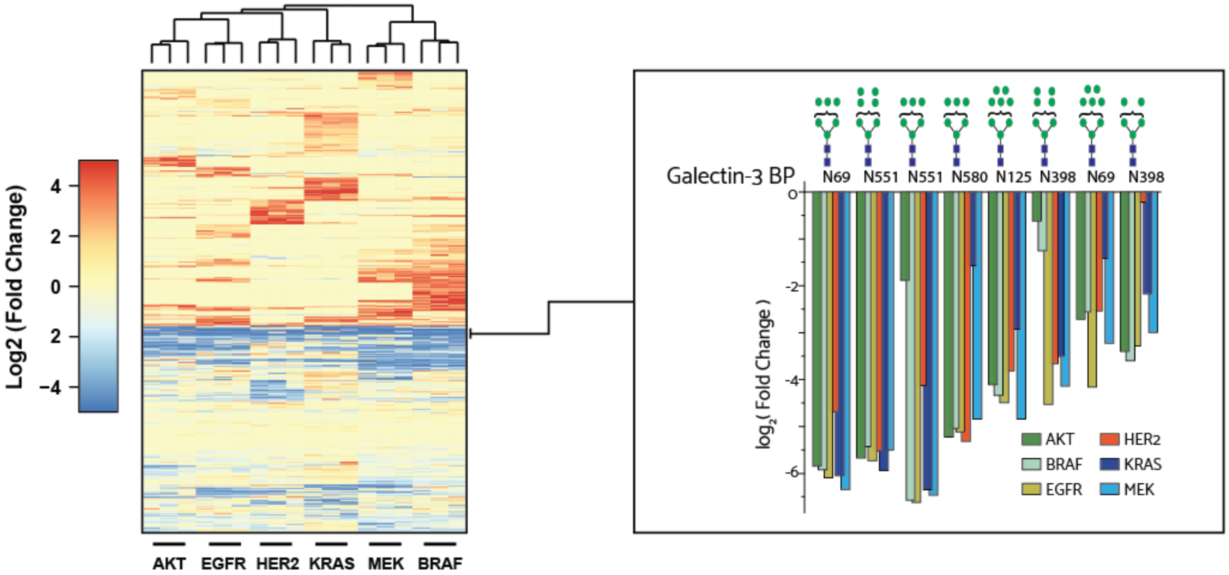


**Figure S14:** A heatmap of glycopeptide fold change compared to empty vector control organized by hierarchical clustering of biological replicates and glycopeptides. A blowout to the right provides the expression of glycopeptides identified from galectin-3-binding protein that constitute a large portion of those species that are similarly down regulated across all cell lines.

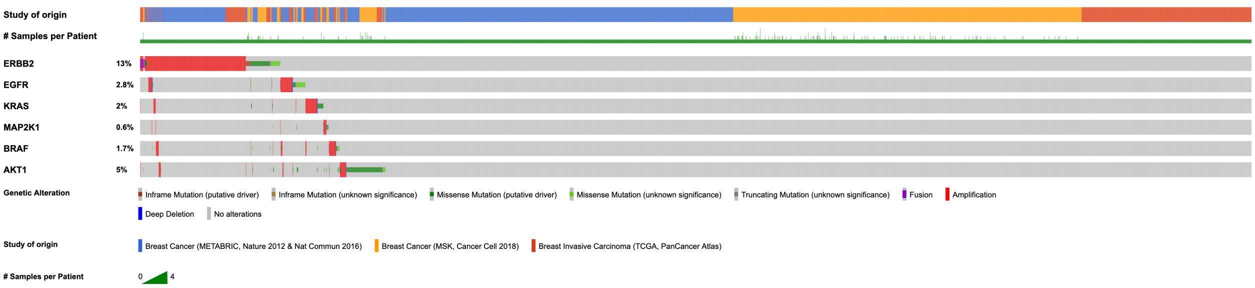


**Figure S15.** Gene alteration in Erbb2, also known as HER2, is more common than KRAS, EGFR, ATK1, BRAF, and MEK in breast cancer. Oncoprint was obtained from querying the six gene of interest in 5349 patient samples across three breast cancer studies (8–10).

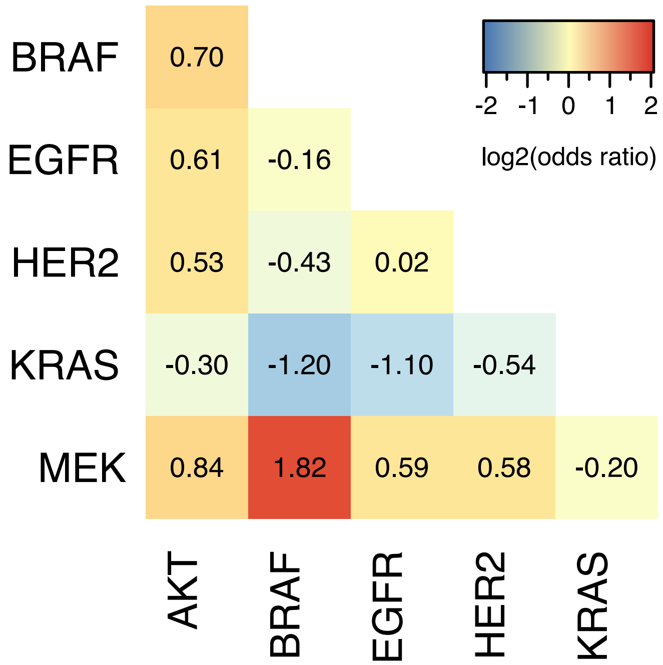


**Figure S16:** An extended in silico comparison of oncogenic mutational occurrence across cancer types indicate a strong mutual exclusivity between BRAF, EGFR, and KRAS. $Log_2$(odds ratio) was obtained from querying the six gene of interest from 20762 patient samples in 33 studies comprising of TCGA PanCancer Atlas Studies (TCGA Research Network: https://www.cancer.gov/tcga) and MSK-IPACT Clinical Sequencing Cohort (11) obtained from cbioportal (12, 13).
